# The Lipidome of iPSC-Derived Retinal Organoids and RPE Partially Resembles that of the Human Retina

**DOI:** 10.64898/2026.06.09.730899

**Authors:** Daniëlle Swinkels, Edwin M. van Oosten, Manon Bouckaert, Anita D.M. Hoogendoorn, Wietske Kieboom, Femke Bukkems, Elfride De Baere, Seba Almedawar, Rob W.J. Collin, Frauke Coppieters, Michèl A.A.P. Willemsen, Frédéric M. Vaz, Alejandro Garanto

## Abstract

New approach methodologies (NAMs), including induced pluripotent stem cell (iPSC)-derived retinal organoids (ROs) and retinal pigment epithelium (iRPE), are increasingly applied to study retinal disease mechanisms and therapeutic strategies. However, these models often remain relatively immature. Given the high lipid content and complex metabolism of the retina, it is unclear to what extent iPSC-derived systems recapitulate the human retinal lipidome. Here, we compared the lipidomic profiles of ROs and iRPE, collected at several differentiation stages, with those of *post-mortem* adult human macular, non-macular and RPE plus choroid (pmRPE). The lipidome of iRPE differed markedly from pmRPE, whereas prolonged differentiation of ROs resulted in a lipidomic profile increasingly resembling that of the *post-mortem* retina. Moreover, ROs showed similarities to both macular and non-macular lipidome. These findings show that iPSC-derived models can become valuable NAMs to study lipid-related retinal disorders and provide a framework to optimize differentiation protocols.

## 1. Introduction

The retina is a highly specialized neural tissue with an exceptionally high lipid content and a complex lipid metabolism (1). There is a continuous renewal of lipid-rich photoreceptor outer segments (POS) as a part of them is phagocytosed daily by the retinal pigment epithelium (RPE). Furthermore, lipids play essential roles in POS biogenesis, phototransduction, synaptic signalling, membrane fluidity, neuroprotection and the visual cycle (2). Consequently, disturbances in lipid synthesis, trafficking or recycling are tightly linked to retinal diseases, ranging from rare inherited metabolic disorders to more common multifactorial conditions such as age-related macular degeneration (2).

Despite the clear importance of lipids for retinal health, our understanding of the lipid composition and functions across distinct retinal cell types remains incomplete. This is partly due to the structural complexity of the retina and the close metabolic coupling between photoreceptors, Müller glia, and the RPE (1, 2). Although murine models have provided important insights into retinal lipid metabolism, species-specific differences in retinal architecture, differences in the diet and lipid-handling pathways limit a direct translatability to the human retina, emphasizing the need for complementary human-based model systems (3–5).

In recent years, induced pluripotent stem cell (iPSC) technology has enabled the generation of human-derived *in vitro* models that retain an individual’s genetic background (6). In the retinal field, iPSC-derived retinal organoids (ROs) and RPE cells (iRPE) have emerged as powerful new approach methodologies (NAMs) to study human retinal development and disease (7–9). ROs self-organize into laminated 3D tissues containing the major retinal cell types and recapitulate key aspects of retinal morphogenesis, cell differentiation and POS formation (8). Accordingly, these models are widely applied to investigate retinal disorders and therapeutic strategies. In parallel, iRPE cells are commonly used to study disease mechanisms, as they reach a high degree of maturation and polarization and are assumed to closely resemble native human RPE (9). In addition to widely used bulk and single-cell transcriptomics, omics analyses such as proteomics and metabolomics are emerging to study disease mechanisms in ROs and iRPE. However, a comprehensive lipidomic profiling of iPSC-derived retinal cell models remains largely unexplored. This could reveal to what extent the lipid composition of Ros and iRPE mirrors that of the human retina, and identify lipids that could help the differentiation, maturation or aging to create better NAMs for disease modelling and therapy development.

Here, to our knowledge, we report for the first time a systematic comparison of the lipidomic profiles of iPSC-derived ROs and iRPE with *post-mortem* human macular, non-macular and RPE. Our results show that iRPE cells do not fully recapitulate the lipidome of *post-mortem* RPE (pmRPE), whereas ROs displayed a lipidomic profile that more closely resembles the *post-mortem* retina and incorporates features of both macular and non-macular regions. These findings provide novel insights to understanding the biology of retinal *in vitro* models, and further improve the differentiation, maturation or ageing to develop relevant NAMs.

## 2. Results

In a recent proof-of-concept study, we showed that reliable lipidomic analysis of ROs requires pooling of six organoids. However, comparisons to previously published retinal lipidomic datasets of the human retina showed clear differences between ROs and their human counterpart, although direct comparison was hampered by methodological differences (Swinkels *et al.* book chapter Retinal Degeneration, under review and (10–13)). These observations prompted us to further expand our analysis and obtain a more comprehensive overview. To this end, we here performed a systematic comparison of the lipidomic profiles of iPSC-derived ROs and RPE with *post-mortem* human macular, non-macular and RPE.

### 2.1. Post-mortem samples characteristics

*Post-mortem* retinal tissue from six subjects were received (three males and three females). Five of those samples were included in the study, of which three males and two females (Table S1). A sixth sample (female) was excluded from the study as the donor was diagnosed with type 2 diabetes, which could alter the lipid content (14, 15). The median donor age was 68 years (mean SD: 69.6±10.1 years) and median *post-mortem* delay was 20 h (mean SD: 17.3±5.8 h).

### 2.2. iRPE have a distinct lipidome compared to post-mortem RPE

We started by comparing the lipidomic composition of iRPE and *post-mortem* RPE. To this end, we performed a principal component analysis (PCA) on the total lipid classes. PCA revealed a clear separation between iRPE and pmRPE samples, indicating that the lipidome of iRPE is fundamentally distinct from that of pmRPE (Fig. 1A).

**Figure 1.**
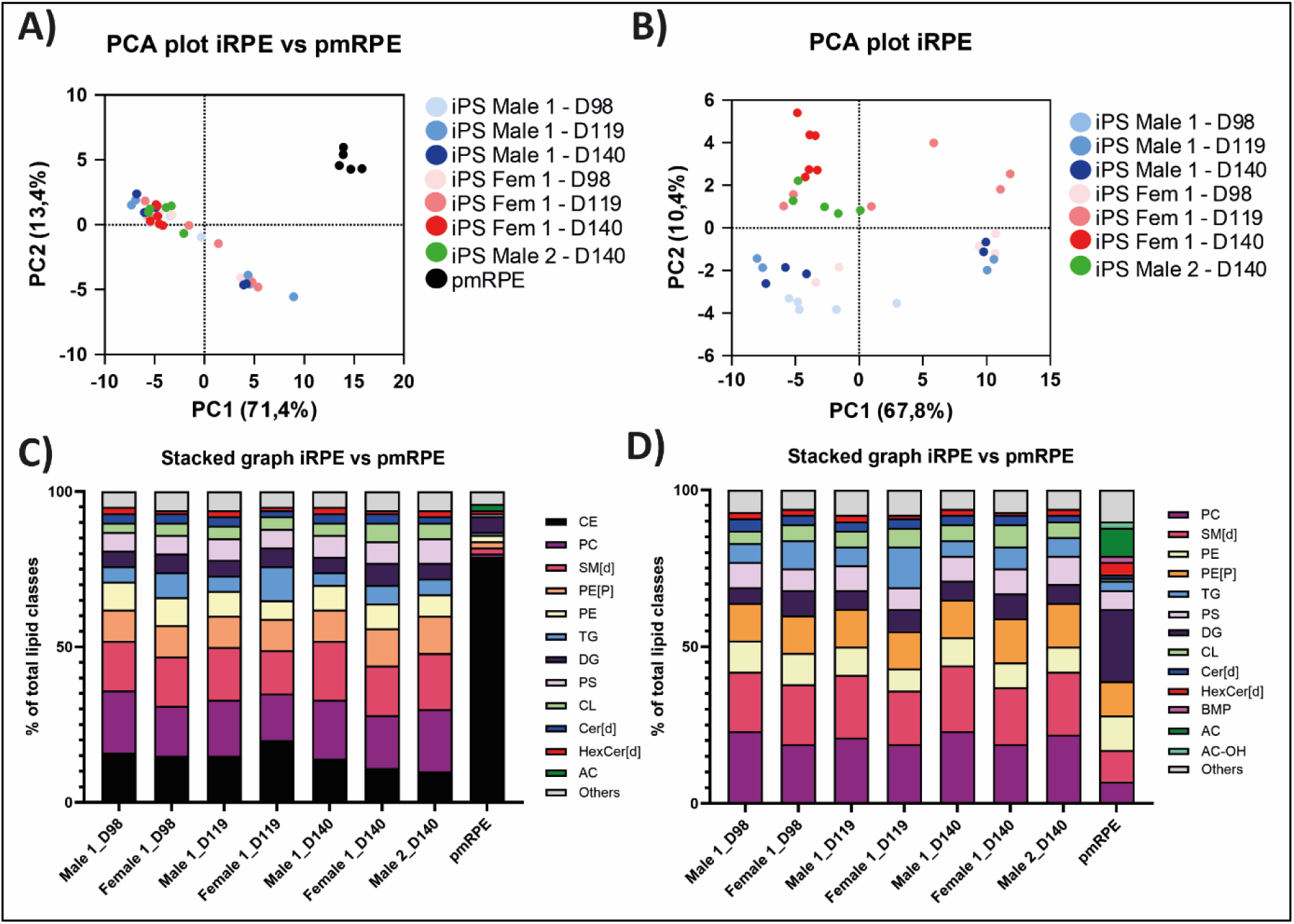
iRPE have a distinct lipidome compared to pmRPE. a) PCA plot showing replicates from pmRPE and different time-points and control lines of iRPE clustered according to their variance in two dimensions. b) PCA plot as in (a) without pmRPE samples. c-d) The relative contribution of different lipid classes to the total lipidome in iRPE and pmRPE with (c) or without CE species (d). AC-acylcarnitines; AC-OH-hydroxy acylcarnitine; BMP-bis(monoacylglycerol)phosphate; CE-cholesteryl esters; Cer[d]-ceramide; CL-cardiolipin; D-differentiation day; DG-diacylglycerol; HexCer[d]-hexosylceramidesPC-phospatidylcholine; PE[P]-alkylphosphatidylethanolamine; PE-phosphatidylethanolamine; PS-phosphatidylserine; SM[d]-sphingomyelin; TG-triglycerides.

Next, we assessed the relative contribution of different lipid classes to the total lipidome in both RPE sample types. This revealed that cholesteryl esters (CEs) accounted for ∼80% of the total lipids measured in pmRPE samples (Fig. 1C). This pronounced enrichment limited direct comparison of the remaining lipid classes and therefore CE species were excluded from subsequent analyses (Fig. 1D). After exclusion of CE species, the relative contribution of lipid classes became more comparable between iRPE and pmRPE, although marked differences remained. While iRPE was characterized by a high abundance of structural phospholipids, particularly phosphatidylcholine (PC; 19–23%) and sphingomyelin [SM(d); 17–21%], the pmRPE lipidome showed a pronounced shift toward neutral lipids. In pmRPE, diacylglycerols (DG; 23%) emerged as the most abundant class, while PC and SM(d) fractions decreased substantially to 7% and 10%, respectively. Crucially, pmRPE contained notable fractions of specific lipids, including acylcarnitines (AC; 9%), hydroxy acylcarnitines (AC-OH; 2%), and bis(monoacylglycerol)phosphate (BMP; 2%), that were not detected in the iRPE model (Fig. 1D). Overall, the lipidome of iPSC-derived RPE is significantly different from *post-mortem* RPE. Of note, the substantial enrichment of CE species was most likely not due to the long time-of-collection period, as samples harvested 10-12h *post-mortem* (pmRPE1,2) showed similar CE levels (Fig. S1A) to those that were collected 20-22h *post-mortem* (pmRPE3-5).

To further evaluate similarities and differences at the biochemical level, we took a closer look into the lipid species within the RPE samples. Fold-change analysis in pmRPE versus iRPE (D140, male 1) revealed that ∼1100 lipid species are >5-fold higher and ∼200 are >5-fold lower (Fig. S1B). A similar trend was observed when comparing pmRPE versus the other iPSC-derived RPE lines at D140 (Table S2). Mainly the more complex lipid species (i.e., containing more than C22 (lysophospholipids), C40 (phospholipid species, C40 designates the sum of the number of carbon atoms in the two radyl units (= side chains)) or C60 (triglycerides, summed number of carbon atoms in three radyl units), or more than 5 double bonds) were enriched in pmRPE. In particular, CE(22:6) and FA(22:6) were amongst the most significantly enriched lipids (Fig. S1B, orange arrows). Despite these pronounced differences in lipid class distribution and lipid species, a substantial overlap was observed at the most abundant species level. Specifically, 22 of the 50 (44%) most abundant lipid species detected in pmRPE (average) were also found among the 50 most abundant species in iRPE (D140) (Table 1). Furthermore, when looking at individual pmRPE samples, this number increased to 26 of the 50 (52%) (Table S3). This indicates that, although the overall lipidome organization differs between the two RPE sources, a core set of highly abundant lipid species is shared.

**Table 1.**
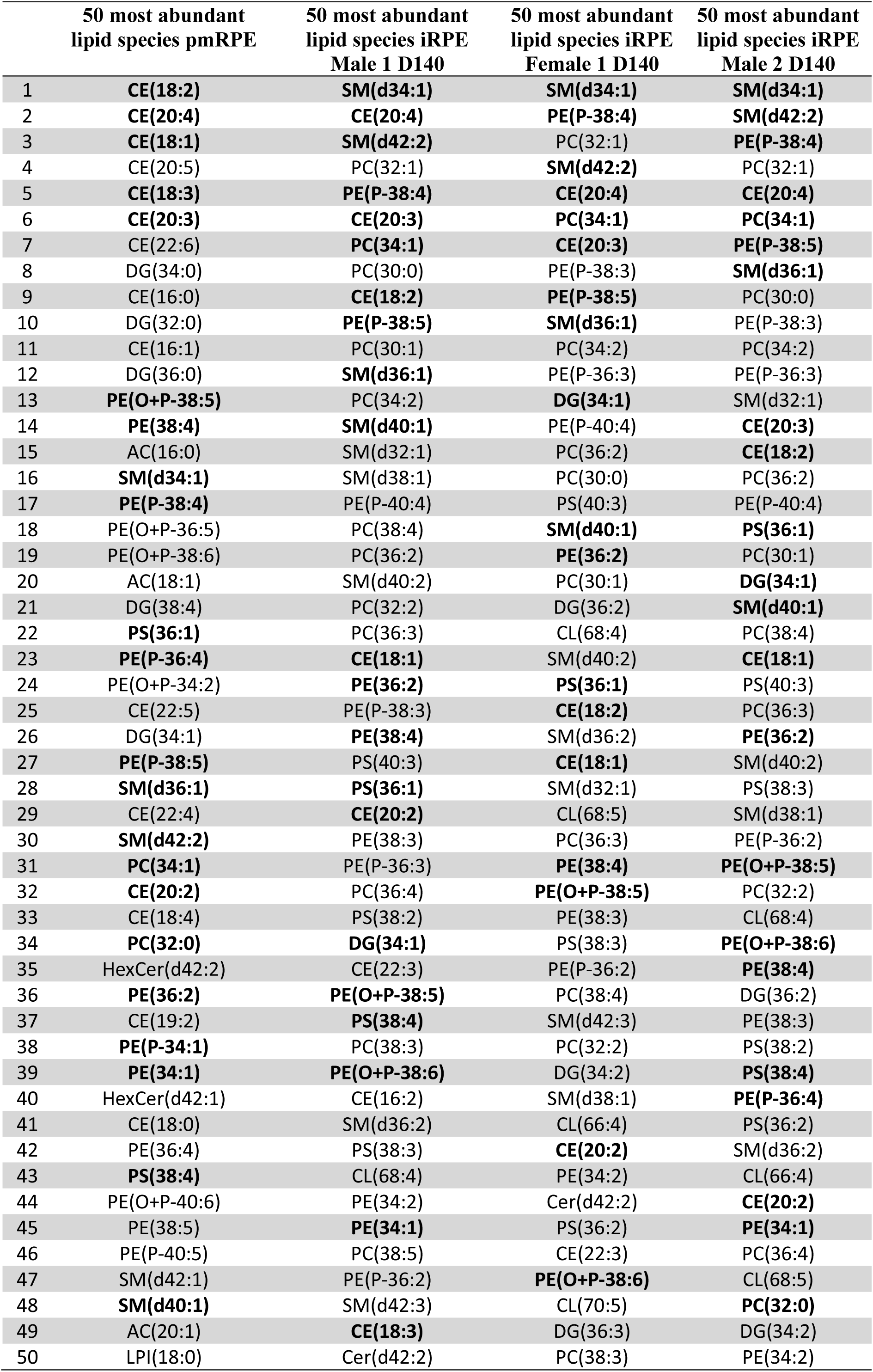
List of the 50 most abundant lipid species in pmRPE and the three different iRPE lines. The common lipid species between pmRPE and iRPE are marked in bold. AC-acylcarnitines; CE-cholesteryl esters; Cer-ceramide; CL-cardiolipin; D-differentiation day; DG-diacylglycerol; HexCer[d]-hexosylceramides; iRPE-iPSC-derived retinal pigment epithelium; LPI-lysophosphatidylinositol; PC-phospatidylcholine; PE[O]-ether phosphatidylethanolamine; PE[P]-alkylphosphatidylethanolamine; PE-phosphatidylethanolamine; pmRPE-post-mortem retinal pigment epithelium; PS-phosphatidylserine; SM[d]-sphingomyelin.

|  | 50 most abundant<br>lipid species pmRPE | 50 most abundant<br>lipid species iRPE<br>Male 1 D140 | 50 most abundant<br>lipid species iRPE<br>Female 1 D140 | 50 most abundant<br>lipid species iRPE<br>Male 2 D140 |
| --- | --- | --- | --- | --- |
| 1 | **CE(18:2)** | **SM(d34:1)** | **SM(d34:1)** | **SM(d34:1)** |
| 2 | **CE(20:4)** | **CE(20:4)** | **PE(P-38:4)** | **SM(d42:2)** |
| 3 | **CE(18:1)** | **SM(d42:2)** | PC(32:1) | **PE(P-38:4)** |
| 4 | CE(20:5) | PC(32:1) | **SM(d42:2)** | PC(32:1) |
| 5 | **CE(18:3)** | **PE(P-38:4)** | **CE(20:4)** | **CE(20:4)** |
| 6 | **CE(20:3)** | **CE(20:3)** | **PC(34:1)** | **PC(34:1)** |
| 7 | CE(22:6) | **PC(34:1)** | **CE(20:3)** | **PE(P-38:5)** |
| 8 | DG(34:0) | PC(30:0) | PE(P-38:3) | **SM(d36:1)** |
| 9 | CE(16:0) | **CE(18:2)** | **PE(P-38:5)** | PC(30:0) |
| 10 | DG(32:0) | **PE(P-38:5)** | **SM(d36:1)** | PE(P-38:3) |
| 11 | CE(16:1) | PC(30:1) | PC(34:2) | PC(34:2) |
| 12 | DG(36:0) | **SM(d36:1)** | PE(P-36:3) | PE(P-36:3) |
| 13 | **PE(O+P-38:5)** | PC(34:2) | **DG(34:1)** | SM(d32:1) |
| 14 | **PE(38:4)** | **SM(d40:1)** | PE(P-40:4) | **CE(20:3)** |
| 15 | AC(16:0) | SM(d32:1) | PC(36:2) | **CE(18:2)** |
| 16 | **SM(d34:1)** | SM(d38:1) | PC(30:0) | PC(36:2) |
| 17 | **PE(P-38:4)** | PE(P-40:4) | PS(40:3) | PE(P-40:4) |
| 18 | PE(O+P-36:5) | PC(38:4) | **SM(d40:1)** | **PS(36:1)** |
| 19 | PE(O+P-38:6) | PC(36:2) | **PE(36:2)** | PC(30:1) |
| 20 | AC(18:1) | SM(d40:2) | PC(30:1) | **DG(34:1)** |
| 21 | DG(38:4) | PC(32:2) | DG(36:2) | **SM(d40:1)** |
| 22 | **PS(36:1)** | PC(36:3) | CL(68:4) | PC(38:4) |
| 23 | **PE(P-36:4)** | **CE(18:1)** | SM(d40:2) | **CE(18:1)** |
| 24 | PE(O+P-34:2) | **PE(36:2)** | **PS(36:1)** | PS(40:3) |
| 25 | CE(22:5) | PE(P-38:3) | **CE(18:2)** | PC(36:3) |
| 26 | DG(34:1) | **PE(38:4)** | SM(d36:2) | **PE(36:2)** |
| 27 | **PE(P-38:5)** | PS(40:3) | **CE(18:1)** | SM(d40:2) |
| 28 | **SM(d36:1)** | **PS(36:1)** | SM(d32:1) | PS(38:3) |
| 29 | CE(22:4) | **CE(20:2)** | CL(68:5) | SM(d38:1) |
| 30 | **SM(d42:2)** | PE(38:3) | PC(36:3) | PE(P-36:2) |
| 31 | **PC(34:1)** | PE(P-36:3) | **PE(38:4)** | **PE(O+P-38:5)** |
| 32 | **CE(20:2)** | PC(36:4) | **PE(O+P-38:5)** | PC(32:2) |
| 33 | CE(18:4) | PS(38:2) | PE(38:3) | CL(68:4) |
| 34 | **PC(32:0)** | **DG(34:1)** | PS(38:3) | **PE(O+P-38:6)** |
| 35 | HexCer(d42:2) | CE(22:3) | PE(P-36:2) | **PE(38:4)** |
| 36 | **PE(36:2)** | **PE(O+P-38:5)** | PC(38:4) | DG(36:2) |
| 37 | CE(19:2) | **PS(38:4)** | SM(d42:3) | PE(38:3) |
| 38 | **PE(P-34:1)** | PC(38:3) | PC(32:2) | PS(38:2) |
| 39 | **PE(34:1)** | **PE(O+P-38:6)** | DG(34:2) | **PS(38:4)** |
| 40 | HexCer(d42:1) | CE(16:2) | SM(d38:1) | **PE(P-36:4)** |
| 41 | CE(18:0) | SM(d36:2) | CL(66:4) | PS(36:2) |
| 42 | PE(36:4) | PS(38:3) | **CE(20:2)** | SM(d36:2) |
| 43 | **PS(38:4)** | CL(68:4) | PE(34:2) | CL(66:4) |
| 44 | PE(O+P-40:6) | PE(34:2) | Cer(d42:2) | **CE(20:2)** |
| 45 | PE(38:5) | **PE(34:1)** | PS(36:2) | **PE(34:1)** |
| 46 | PE(P-40:5) | PC(38:5) | CE(22:3) | PC(36:4) |
| 47 | SM(d42:1) | PE(P-36:2) | **PE(O+P-38:6)** | CL(68:5) |
| 48 | **SM(d40:1)** | SM(d42:3) | CL(70:5) | **PC(32:0)** |
| 49 | AC(20:1) | **CE(18:3)** | DG(36:3) | DG(34:2) |
| 50 | LPI(18:0) | Cer(d42:2) | PC(38:3) | PE(34:2) |

### 2.2. Lipid content in iRPE does not fluctuate much during maturation

Finally, it was also of interest to investigate whether the lipidome of iRPE changes during maturation. Therefore, samples were collected at D98, D119, and/or D140. PCA and lipid class analyses did not reveal substantial differences between these timepoints (Fig. 1B, D). Instead, variability appeared to be primarily driven by inter-line differences rather than differentiation time. This was also supported by the hierarchical clustering dendrogram, which did not show differences between clustering of the D98, D119 nor D140 iRPE to the pmRPE samples (Fig. S1C). Overall, these data suggest that the lipidome of iRPE reaches a relatively stable lipid state by D98 under the conditions examined.

### 2.3. Feeding POS to iRPE does not sufficiently modify the lipid profile to match that of pmRPE

In the retina, under physiological conditions, approximately 10% of the POS are shed daily and subsequently phagocytosed by the RPE. The lipid content of POS accounts to nearly 60% of their total composition. In addition, the RPE is in direct contact with the systemic circulation and is therefore exposed to lipids from hepatic metabolism (2). Both sources contribute substantially to the lipid homeostasis of the RPE and are absent in the iRPE *in vitro* model system. We therefore investigated whether supplementation of iRPE with porcine POS could alter the lipidomic profile to make it more comparable to that of pmRPE. To this end, iRPE were exposed to porcine POS for five days. As a negative control condition, iRPE cells were similarly treated with only FCS, which at the same time is a source of lipids, as iRPE are cultured without FCS.

Overall, POS treatment resulted in >5-fold higher levels of ∼800 lipid species and >5-fold lower levels of ∼129 lipid species compared to untreated iRPE samples (Fig. S2B and Table S4). However, despite POS exposure, iRPE samples continued to cluster distinctly from pmRPE samples (Fig. 2A). In line with this observation, the overall distribution of lipid classes in POS-treated iRPE remained markedly different from that observed in pmRPE (Fig. 2C), also after exclusion of the aforementioned CE species (Fig. 2D). Furthermore, ∼1000 lipid species were either significantly higher or lower in pmRPE compared to POS-treated iRPE (Fig. S2C). In addition, the untreated iRPE cells formed a distinct cluster from FCS– and POS-treated iRPE, highlighting the contribution of both FCS and POS to the lipidome (Fig. 2B and Fig. S2A).

**Figure 2.**
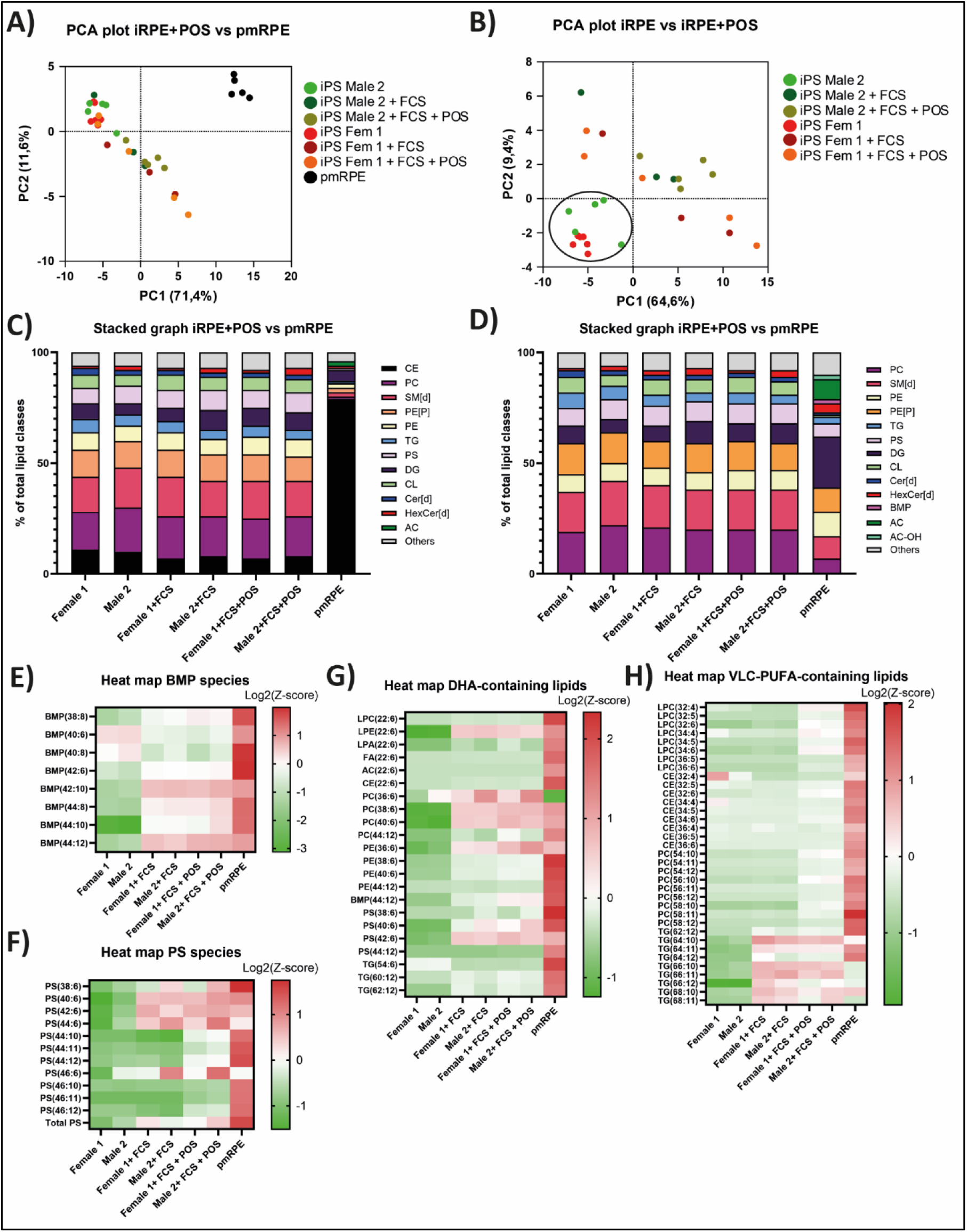
Feeding POS to iRPE does not sufficiently modify the lipid profile to match that of pmRPE. a) PCA plot showing replicates from pmRPE and different time-points and control lines of iRPE treated with POS (photoreceptor outer segments) clustered according to their variance in two dimensions. b) PCA plot (as in panel a) without pmRPE samples. Black circle highlights untreated iRPE samples. c-d) The relative contribution of different lipid classes to the total lipidome in pmRPE and iRPE with (c) or without CE species (d). e-h) Heat map of BMP (e), PS (f), DHA-containing (g) and VLC-PUFA-containing (f) lipids. Data are represented as log2(FC). AC-acylcarnitines; AC-OH-hydroxy acylcarnitine; BMP-bis(monoacylglycerol)phosphate; CE-cholesteryl esters; Cer[d]-ceramide; CL-cardiolipin; D-differentiation day; DG-diacylglycerol; HexCer[d]-hexosylceramides; PC-phospatidylcholine; PE[P]-alkylphosphatidylethanolamine; PE-phosphatidylethanolamine; PS-phosphatidylserine; SM[d]-sphingomyelin; TG-triglycerides.

To investigate this further, we took a closer look into specific lipid species and classes related to POS phagocytosis. Firstly, BMPs are specialized, negatively charged phospholipids enriched in late endosomes and lysosomes in RPE cells. During POS phagocytosis, BMP functions as a docking platform recruiting positively charged lipid hydrolases and facilitating lipid degradation (16, 17). Indeed, specific BMP species, most likely containing one docosahexaenoic acid (DHA, C22:6) and one unsaturated fatty acid, were upregulated upon FCS and POS treatment compared to untreated iRPE cells (Fig. 2E and Fig. S2B). Interestingly, POS treatment, but not FCS treatment, increased levels of some BMP species (e.g., BMP(44:12), most likely 2 x C22:6/DHA-containing) to levels of the pmRPE (Fig. 2E). Secondly, PS is normally confined to the cytoplasmic leaflet of the POS membranes, but upon light exposure flips to the extracellular surface, where it serves as a critical “eat-me” signal, thereby initiating recognition and binding of POS by the RPEs (17, 18). Therefore, an increase in PS species can be expected in the POS-treated samples. However, PS species increased in both FCS and POS-treated samples, compared to untreated iRPE cells (Fig. 2F). However, total PS levels and PS species most likely containing the retina-specific lipids DHA and very-long-chain polyunsaturated fatty acids (VLC-PUFAs) were higher in POS-treated samples, compared to FCS-treated samples (Fig. 2F and Fig. S2D). Lastly, we investigated lipid species that are enriched in POS *in vivo*. POS are mainly composed of PC and PE species (∼80% of the total lipids) enriched in DHA and VLC-PUFAs (19). Therefore, we next examined the effect of FCS and POS supplementation on these lipid species. Indeed, levels of DHA-and VLC-PUFA–containing lipids were increased in both supplementation methods and, in some cases, approached those observed in pmRPE (Fig. 2G-H). However, it seems that these levels are primarily driven by FCS supplementation rather than by POS exposure. Together, these findings indicate that while supplementation with FCS and POS can partially modulate specific lipid classes in iRPE, it is insufficient to fully recapitulate the lipidomic profile of pmRPE under the tested conditions.

### 2.4. The lipid profile of the human post-mortem macular is distinct from the non-macular

Agbaga *et al.* previously showed that the lipid composition of rods and cones differs, by studying cone-and rod-dominant animal retinas (20). As the macular is enriched in cone photoreceptors, whereas the peripheral retina is dominated by rods, this analysis is particularly relevant in the context of retinal organoids, which predominantly contain cones. Therefore, we here, for the first time, investigate the lipidomic profile of *post-mortem* human macular versus non-macular regions from the same individual, to assess whether regional differences in photoreceptor composition are reflected in the retinal lipidome.

PCA revealed complete separation between macular and peripheral (non-macular) retinal samples, with each group forming a distinct, non-overlapping cluster (Fig. 3A). This segregation underscores substantial regional differences in the lipidomic architecture of cone-rich versus rod-rich retinal areas. Evaluation of relative lipid class distributions further emphasized these disparities (Fig. 3B). The macular lipidome was primarily characterized by an abundance of structural phospholipids, specifically PC (20%), PE[P] (14%), SM[d] (13%), and CL (11%). In contrast, the peripheral retina exhibited a marked shift toward neutral lipid accumulation; DG emerged as the dominant class (20%), accompanied by substantial depletions in both PC (9%) and SM[d] (5%).

**Figure 3.**
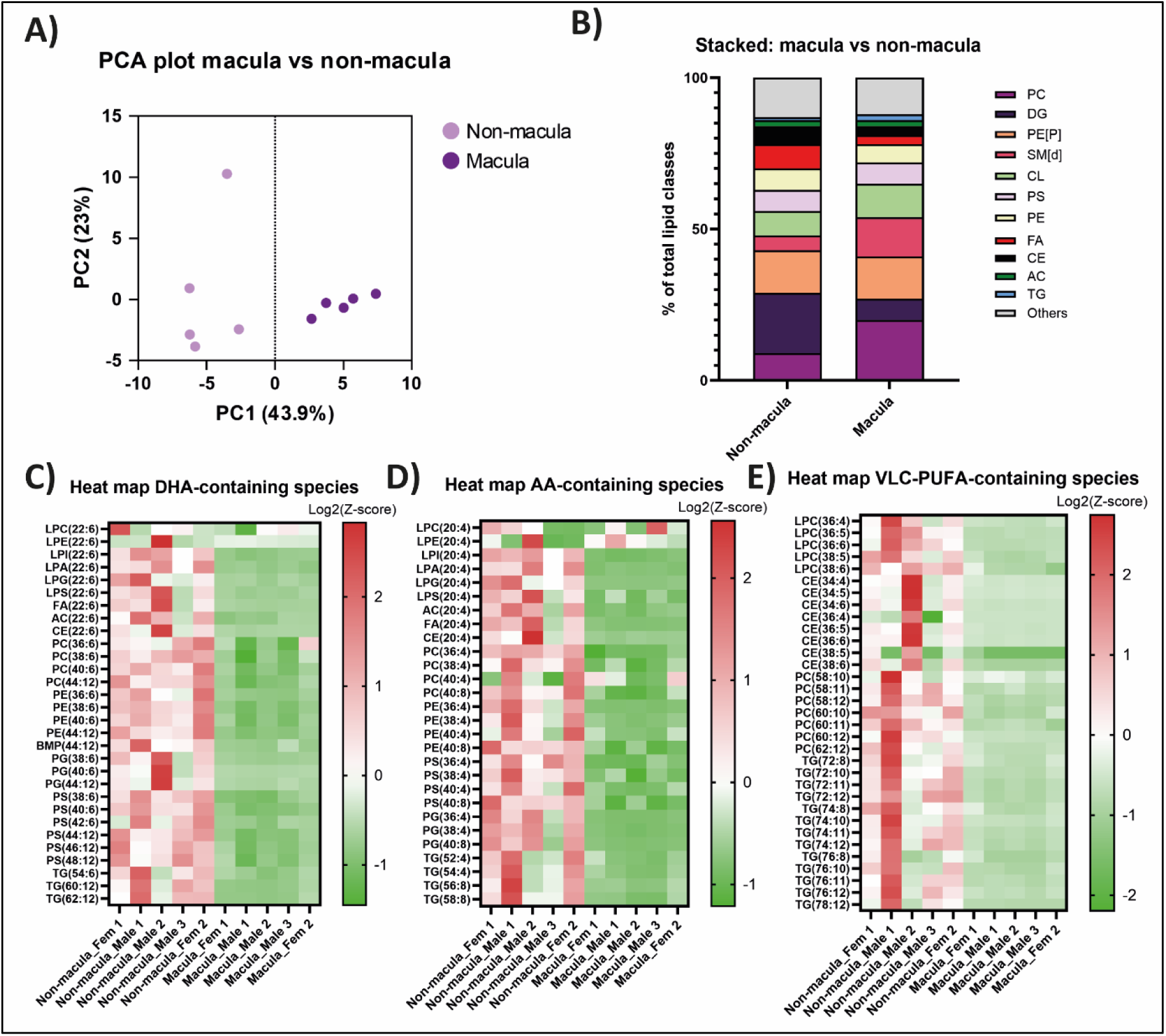
A distinct lipid profile in post-mortem macular versus non-macular. a) PCA plot showing replicates from post-mortem macular and non-macular samples clustered according to their variance in two dimensions. b) The relative contribution of different lipid classes to the total lipidome in macular and non-macular samples. c-e) Heat map of DHA-(c), AA-(d), and VLC-PUFA-(e) containing lipid species. Data are represented as log2(FC). (L)PC-(lyso)phospatidylcholine; (L)PE-(lyso) phosphatidylethanolamine; (L)PG-(lyso)phosphatidylglycerol;(L)PI-(lyso)phosphatidylinositol; (L)PS-(lyso)phosphatidylserine; AC-acylcarnitines; BMP-bis(monoacylglycerol)phosphate; CE-cholesteryl esters; CL-cardiolipin; DG-diacylglycerol; FA-fatty acid; PE[P]-alkylphosphatidylethanolamine; SM[d]-sphingomyelin; TG-triglycerides.

Given the known importance of PUFAs in photoreceptor function and survival, we next focused on DHA, VLC-PUFAs and arachidonic acid (AA)-containing lipid species. Previous studies in animal models showed that these lipid species are more abundant in rods than cones (20). To determine whether this regional enrichment is conserved in the *post-mortem* retina, we examined DHA– and VLC-PUFA-containing lipid species across multiple lipid classes, including (lyso)phospholipids, CEs, BMPs and TGs. Heat map analysis revealed a consistent enrichment of DHA-, AA– and VLC-PUFA-containing species in non-macular samples compared to macular samples (Fig. 3C-E). This pattern indicates that rod-enriched regions of the *post-mortem* retina harbour higher levels of these lipid species than cone-rich regions, which is consistent with observations from animal models (20).

To further investigate the differences between the macular and non-macular lipidome, we investigated the fold-change of lipid species. Overall, 429 lipid species were >5-fold higher and 47 were >5-fold lower in macular samples than non-macular (Fig. S3A). The main differences could be found in the PC, TG, TG[O], PC[O] and PC[P] species. Mainly the simpler lipid species (i.e., containing less than C20 (lysophospholipids), C40 (phospholipid species) or C60 (triglycerides), or less than 5 double bonds) were enriched in macular samples (Fig. S3A and Table S5).

### 2.5. The lipidome of ROs changes throughout the differentiation process

Next, we investigated if the lipidome differs throughout the differentiation process of retinal organoids. Therefore, samples were collected at D140 (no brush border), D180 (brush border, mainly cones and few rods) and D240 (brush border, cones and more rods). Interestingly, most of the D240 samples cluster separately from the D140 and D180 ROs (Fig. 4A and Fig. S3B). Of note, although there is still inter-line variability, it is less pronounced in the ROs samples compared to the iRPE samples (Fig. 4A, 1B). Subsequently, we took a closer look into the difference between the D140 and D180 samples versus the D240 samples. Firstly, this could be attributed to a differential distribution of lipid classes, where D140 and D180 samples have a higher content of CE and TG species, and lower levels of DG, SM[d] and AC species, compared to D240 samples (Fig. 4B). Also, a clear difference in AA– and VLC-PUFA-containing lipid species was found (Fig. 4C-E). While AA-containing lipid species were enriched in D140 and D180 samples compared to D240 samples, the opposite was true for VLC-PUFA-containing lipid species, most likely to compensate for the degree of unsaturation in the retina. Overall, PC-species up to 62 carbons (e.g., C18 plus C44) and TG-species up to 76 carbons (e.g., 2 times C18 plus C40) could be reliably measured, showcasing that also retina-specific lipid species can be detected in the ROs. In conclusion, the lipidome of ROs changes throughout the differentiation process.

**Figure 4.**
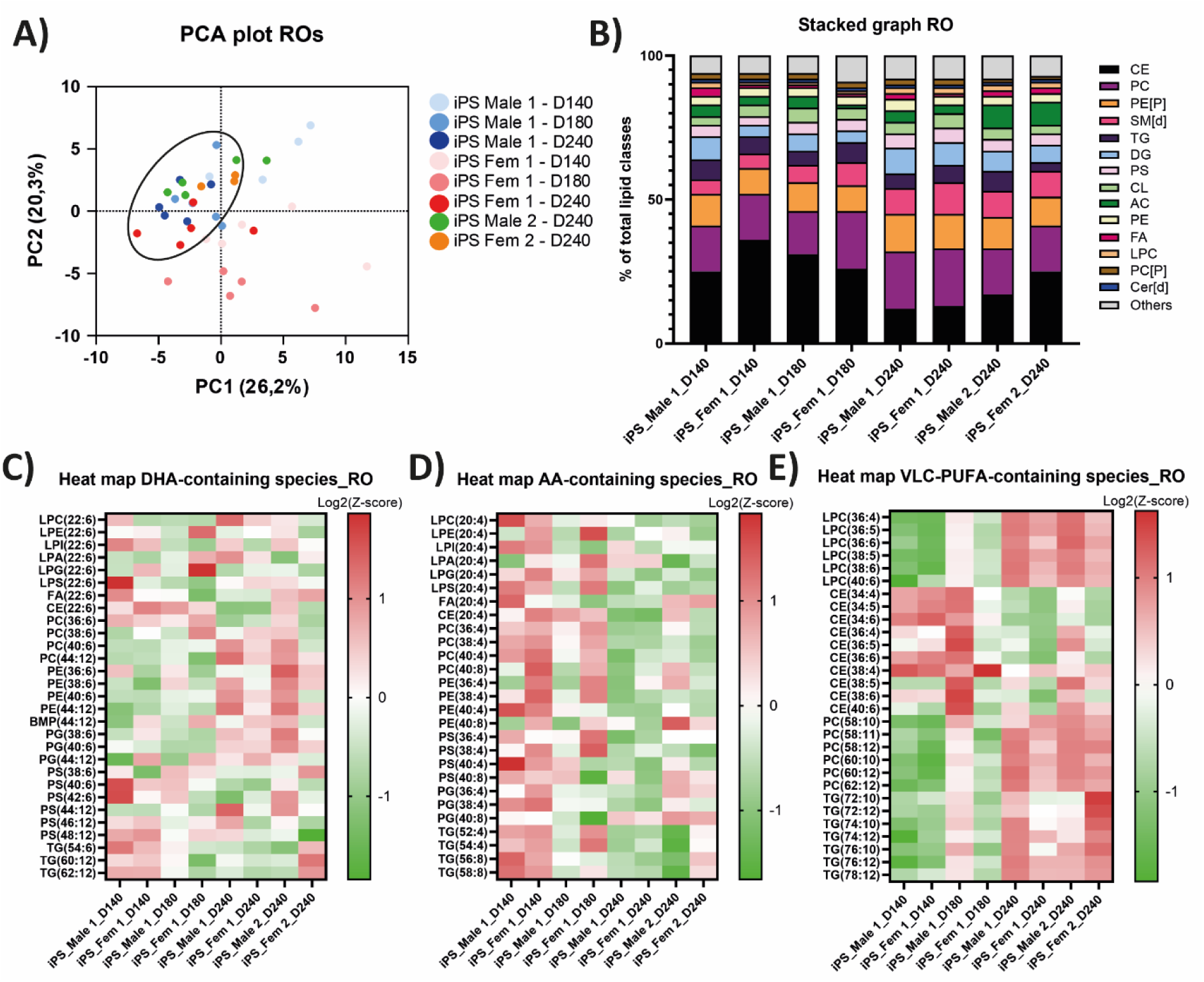
The lipidome of ROs changes throughout the differentiation process. a) PCA plot showing replicates from different time-points and control lines of ROs clustered according to their variance in two dimensions. Black circle highlights D240 samples. b) The relative contribution of different lipid classes to the total lipidome in RO samples. c-e) Heat map of DHA– (c), AA– (d), and VLC-PUFA– (e) containing lipid species. Data are represented as log2(Z-score). (L)PC-(lyso)phospatidylcholine; (L)PE-(lyso) phosphatidylethanolamine; (L)PG-(lyso)phosphatidylglycerol;(L)PI-(lyso)phosphatidylinositol; (L)PS-(lyso)phosphatidylserine; AC-acylcarnitines; CE-cholesteryl esters; CL-cardiolipin; DG-diacylglycerol; FA-fatty acid; PE[P]– alkylphosphatidylethanolamine; SM[d]-sphingomyelin; TG-triglycerides.

### 2.6. The mature RO lipidome resembles features of both the post-mortem macular and non-macular lipidome

Given the difference in lipidomic profile of the *post-mortem* macular versus non-macular, the differential distribution of cones and rods in these regions, and that ROs display a maturation-dependent shift in their lipidome, we next investigated whether and at what differentiation stage the lipidome of the ROs better resembles the macular or non-macular lipidome. However, when plotting the data on a PCA plot, the ROs are clustering separately from the *post-mortem* macular and non-macular samples. (Fig. 5A and Fig. S3B).

**Figure 5.**
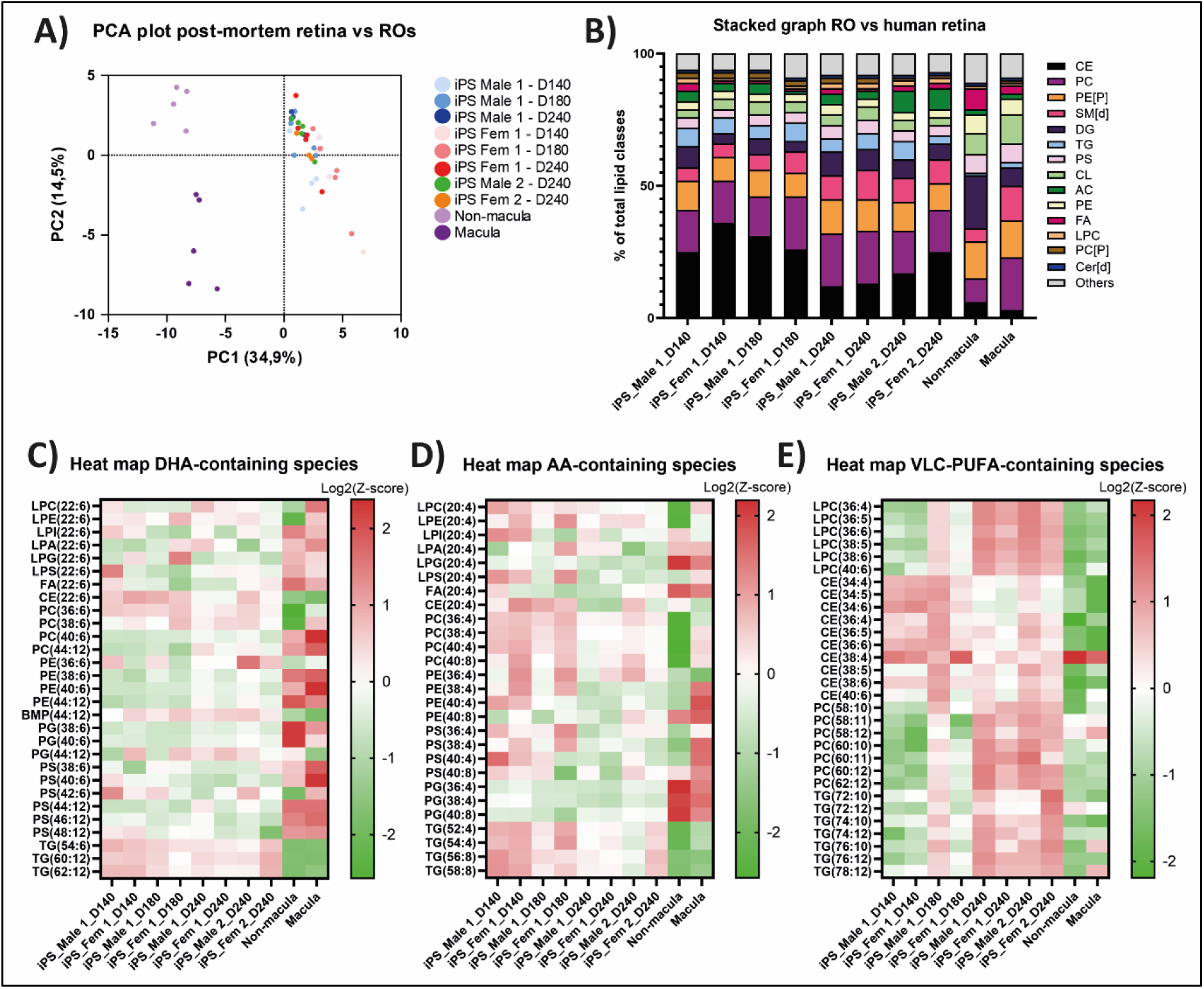
The mature RO lipidome resembles the post-mortem macular more than the non-macular lipidome. a) PCA plot showing replicates from post-mortem macular, non-macular and different time-points and control lines of ROs clustered according to their variance in two dimensions. b) PCA plot as a) without post-mortem samples. Black circle highlights D240 samples. c) The relative contribution of different lipid classes to the total lipidome in post-mortem and RO samples. d) e-g) Heat map of DHA-(e), AA– (f), and VLC-PUFA– (g) containing lipid species. Data is represented as log2(Z-score). (L)PC-(lyso)phospatidylcholine; (L)PE-(lyso) phosphatidylethanolamine; (L)PG-(lyso)phosphatidylglycerol;(L)PI-(lyso)phosphatidylinositol; (L)PS-(lyso)phosphatidylserine; AC-acylcarnitines; CE-cholesteryl esters; CL-cardiolipin; DG-diacylglycerol; FA-fatty acid; PE[P]-alkylphosphatidylethanolamine; SM[d]-sphingomyelin; TG-triglycerides.

Subsequently, we evaluated the differential distribution of lipid classes between ROs and *post-mortem* retina samples (Fig. 5B). Overall, the lipidomic profiles of ROs and *post-mortem* retinas were substantially more comparable than the iRPE and pmRPE models. While both RO and *post-mortem* samples comprised the same major lipid classes, their relative abundances varied. The RO lipidome was characterized primarily by a highly variable abundance of CE (12–36%) and structural phospholipids, including PC (15–20%) and PE[P] (9–13%). This overall distribution exhibited mixed characteristics of the *in vivo* tissue, slightly favoring the macular lipidome while retaining similarities to the peripheral retina. Notably, the lipid class distribution in mature organoids (D240) converged more closely toward both *post-mortem* profiles, particularly regarding the stabilization of CE and SM[d] proportions, compared to earlier developmental stages (D140 and D180). This trajectory underscores a maturation shift in the RO lipidome toward an *in vivo*-like human retinal signature.

We then studied retina-specific lipid species (i.e., AA, VLC-PUFA and DHA) in macular, non-macular and RO samples (Fig. 5C-E). For DHA-containing lipid species, no clear conclusions could be drawn. For some lipid species, levels were higher than *post-mortem* retina samples, but for others lower. For AA-containing lipid species, levels were higher in younger ROs (D140-D180), compared to older ROs (D240), and better resembled the macular lipid levels (Fig. 5D). On the other hand, VLC-PUFA-containing lipids levels were higher in older ROs (D240) compared to younger ROs (D140-D180) and *post-mortem* retina samples (Fig. 5E).

Lastly, we investigated the fold-change of lipid species between macular, non-macular and ROs at D240. Overall, 255 lipid species were >5-fold higher and 1097 were >5-fold lower in non-macular samples than ROs (Fig. S3C). For the macular, 242 species were >5-fold higher and 555 were >5-fold lower than ROs samples (Fig. S3D). The same trend was also observed for the other iPSC-derived ROs (Table S5). This again points towards a slightly better resemblance of the RO lipidome to the macular lipidome.

Overall, it can be concluded that ROs slightly better represent the macular lipidome, although also show similarities with the non-macular lipidome. Importantly, mature ROs (D240) show a more similar lipid profile to the *post-mortem* samples than younger ROs (D140-180), indicating a lipidome shift during maturation.

### 2.7 No clear differences in lipid profile between male and female retina

Despite the low number of samples, we attempted to assess whether biological sex contributed significantly to the lipidome. Therefore, PCA assays were performed on lipidomic profiles from the *post-mortem* retina and pmRPE, and iPSC-derived ROs and RPE samples (at least n=2). Across all tissue types, PCA did not reveal a clear segregation of samples based on sex (Fig. S4). For the iPSC-derived samples (both iRPE and RO) there was a tendency to segregation, which could indicate a possible contribution by biological sex, or could be caused by potential inter-line variability. Overall, these data do not seem to indicate that sex significantly influenced the retinal lipidome when looking at lipid classes under the experimental conditions and sample numbers examined, however further studies and more samples are required to confirm these findings.

## 3. Discussion

In this study, we provide the first systematic comparison of the lipidomic profiles of iRPE cells and ROs with *post-mortem* human macular, non-macular and RPE. Our data revealed pronounced differences between iRPE and pmRPE, whereas ROs displayed a lipidomic profile that more closely resembled the *post-mortem* retina where both macular and non-macular regions were represented. Some explanations for these differences could be associated with the age of the donors, compared to our embryonic retinal models. Still, these data are highly valuable as they can provide novel insights on which compounds to deliver to the medium to for instance improve maturation, but also to age these models and accelerate the onset of phenotypes. Also, by looking at the species represented in the lipidomes of the *in vitro* models, our results suggest that *in vitro* retinal cell models can reliably be used to investigate lipid-related retinal disorders.

An important observation was the variability in lipid distribution between control iPSC-derived RO and RPE lines. This did not seem to be protocol dependent, as the RMCGENi005-A-1-derived RO samples (differentiated using (21)) clustered together with the other control lines derived RO (differentiated using (22)). Therefore, it is believed that this inter-line variability is rather due to differences in the genetic background, tissue of origin or reprogramming method. Overall, this underscores the importance of using isogenic controls when modelling lipid-related retinal diseases.

We also evaluated how lipid composition changes during the differentiation of iRPE and ROs. Interestingly, the lipidome of iRPE remained stable over time, indicating that prolonged culture alone is insufficient to promote maturation towards a more human-like RPE lipid profile. In contrast, ROs exhibited a clear lipid progression, with longer differentiation times resulting in a lipidomic profile that increasingly resembled that of the *post-mortem* retina. These findings support the concept that ROs undergo ongoing metabolic maturation and underscore the importance of extended culture periods for lipid-centred studies. In this study, organoids were differentiated only up to D240. However, these results raise the question whether differentiation to later timepoints (e.g., D280) would yield a lipidome that even more closely mirrors the lipid profile of the *post-mortem* retina.

The lack of concordance between iRPE and pmRPE lipidomes is an essential finding, as these models are an important model to study retinal diseases where RPE is affected, including age-related macular degeneration (23). pmRPE samples were dominated by cholesteryl esters, accounting for approximately 80% of the total lipid content, whereas these species constituted only around 10% in iRPE. This discrepancy may reflect donor age, as cholesteryl esters are known to accumulate in the human RPE over time. In addition, our *post-mortem* RPE samples also contained parts of the choroid and Bruch’s membrane where often accumulation of lipids occur (24–26). These two factors may explain the big differences in the lipid content and why the iRPE models do not fully recapitulate the lipid composition of native *post-mortem* RPE. This limitation is particularly relevant for studying lipid-related and age-associated retinal disorders. It would be interesting to compare pure pmRPE (without choroid and Bruch’s membrane) to the iRPE lipidome. Therefore, FACS-sorting to separate the pmRPE from the choroid and Bruch’s membrane could be a possible alternative.

Other reasons for the discrepancy between the iRPE and the pmRPE lipidome could be the lack of a vascular system and that there is no POS phagocytosis, both of which are critical for lipid transport and homeostasis *in vivo* (2). To partially overcome these limitations, we supplemented porcine POS to the iRPE. Under physiological conditions, each RPE cell ingests approximately 10 POS daily and current protocols describe short-term (i.e., 2-5 hours) POS exposure experiments using 10 POS/RPE cell (27). However, as we wanted to make a more human-like situation but avoiding excessive lipid loading, we opted for a 5-day treatment with 5 POS per iRPE cell. Indeed, POS feeding resulted in higher levels of lipid species and classes that are related to POS phagocytosis, such as BMP, PS and phospholipid species containing DHA and VLC-PUFAs. However, POS feeding did not normalize lipid levels to those of the pmRPE. While it remains possible that a higher POS load would have given different results, we consider this unlikely as we do not expect POS feeding to alter the CE levels in such a short time. Therefore, perhaps longer incubation periods are required to start observing accumulation of lipids associated with aging. Additionally, most changes upon POS feeding were also observed in the FCS-treated condition. Therefore, we propose that either POS feeding should be extended (e.g., 10 POS/RPE cell for 2 weeks or more), or FCS should be replaced with the RPE phagocytic receptor ligand proteins like MFG-E8, GAS6 and Protein S, to see direct contribution of POS (27). However, although this last option would allow a cleaner comparison, FCS could also be seen as the systemic lipid supplementation and therefore an essential supplement for RPE differentiation. Lastly, it would be interesting to see if co-cultures of iRPE and endothelial cells and/or retinal organoids will impact the lipid accumulation in iRPE.

The human retina has been extensively characterized using proteomics and (single-cell) RNA sequencing approaches. However, comprehensive untargeted lipidomic profiling of human retinal tissue remains limited, and existing studies do not provide a complete overview of the retinal lipidome (10–13). Moreover, previous analyses did not distinguish between macular and peripheral regions, despite evidence that this distinction is biologically relevant (20). Agbaga *et al.* demonstrated differences in lipid composition between rods and cones using rod– and cone-dominant animal retinas, suggesting that regional lipid specialization is likely present in the human retina as well. Here, we present the first direct comparison of the lipid composition of *post-mortem* human macular and peripheral retina. We confirmed the findings of Agbaga *et al.* and identified substantial regional differences in their lipidomic profiles. By making this dataset publicly available, we aim to provide a valuable resource that facilitates mechanistic insight and supports therapeutic target discovery in retinal disorders.

Interestingly, the RO lipidome showed a remarkable similarity to that of the human macular and non-macular, supporting the notion that ROs encompass features of both retinal regions. These data are reassuring as it was always believed that the ROs more resemble the macular, as they are cone-rich (8). Another interesting finding was the detection of VLC-PUFAs in ROs, underscoring the capacity of ROs to synthesize complex retinal-specific lipid species. However, clear differences remain between the RO and *post-mortem* retina lipidome, and addition of DG-, CL-, PE-species, and free fatty acids to the media of ROs could resolve this issue. Indeed, West *et al.* showed that DHA supplementation to the RO differentiation medium improved POS formation (28). Therefore, addition of DHA and above-mentioned lipids, could help to further mature these organoids. Moreover, in a recent study we showed that light stimulation improved RO maturation as the number of rods increased (22). Therefore, comparing the two protocols (i.e., lipid supplementation and light exposure) could shed light on whether ROs become even more similar to the human retina in terms of lipid content.

We were unable to robustly examine potential sex-dependent differences in retinal lipid composition due to the low number of samples included in the study. This question is particularly relevant considering recently published reports suggesting an influence of sex on retinal lipid metabolism (29). Future studies with larger cohorts will be required to determine whether such differences are also present in human retinal tissue and iPSC-derived retinal models.

Lastly, several lipid classes were enriched in the iPSC-derived models compared to the *post-mortem* samples. More specifically, structural phospholipids (e.g., PC, SM[d]) exhibited a higher proportion in iRPE and ROs, while neutral lipids (e.g., CE, TG, DG) were higher in *post-mortem* samples. Structural lipids affect membrane rigidity, receptor signalling, vesicle trafficking, and protein-lipid interactions (30). Conversely, lower levels of neutral lipids reduce intracellular energy storage and lipid droplet-mediated signalling (31). These differences could be attributed to the fetal-like metabolic state of iPSC-derived models and the absence of systemic lipid supply, highlighting a key consideration when using these models to recapitulate adult human tissue physiology. Therefore, optimizing the lipid composition of iPSC-derived retinal models is crucial to improve their ability to recapitulate human retina physiology and function.

Overall, our data provide a framework to define specific lipid species that could be supplemented to promote a more physiologically relevant, aged *post-mortem* retina and RPE phenotype. For the iRPE, some possibilities could be to evaluate medium supplemented with LDL-cholesterol, enhancing acyl-coenzyme A cholesterol acyltransferase (ACAT) activity to drive CE availability and esterification (32–34), co-culturing with ROs and endothelial cells, supplementing the media with A2E to support ageing (35, 36), and/or longer POS supplementation using the phagocytic receptor ligand proteins (MFG-E8, GAS6 and Protein S) (27). But first, it would be essential to understand the lipidomic composition of pure pmRPE (without choroid and Bruch’s membrane). For the ROs, more mature models could be established by light exposure (22) or supplementation of certain lipids, such as DG-, CL-, PE-species and free fatty acids, especially DHA (28).

In conclusion, we provide the first direct comparison of the lipid composition of *post-mortem* human macular and non-macular retina and revealed substantial regional differences. Furthermore, we showed that the lipid composition of iRPE is different from the *post-mortem* RPE. ROs, on the other hand, mirrored parts of the macular and non-macular lipid profile and retina-enriched lipid species, such as DHA, AA and VLC-PUFAs, were detectable especially at later differentiation stages. Therefore, *in vitro* retinal models can become robust NAMs to study lipid-driven retinal disease mechanisms and evaluate therapies, although some optimizations are required.

## 4. Materials and Methods

### 4.1. iPSC culturing and differentiation into ROs and iRPE

Control iPSCs were cultured in Essential 8 Flex (E8F, Gibco, A2858501) medium on Geltrex-coated plates (Gibco, A1413302) at 37 °C and 5% CO2. Four control iPSC lines were used; two male-derived (SCTCi048-A-1 (also known as IPS23-00102, unpublished, male 1) and HPSI0114i-kolf_3 (KOLF3, culture collections, male 2)) and two female-derived (SCTCi010-A, female 1 (37) and RMCGENi005-A-1, female 2 (38)). Differentiation of iPSCs into ROs was based on a 2D/3D protocol described elsewhere (39), with some adjustments that were extensively detailed here (22). Differentiation of RMCGENi005-A-1 towards retinal organoids was based on (21), but the maintenance medium from differentiation day (D) 100 until collection was the same as used for the other lines. iPSCs were differentiated into iRPE using the direct differentiation protocol published elsewhere (40). At passage 3, samples were transferred to 24-well inserts (cellQart, 9320412) or 6-well inserts (cellQart, 9300412).

Four control iPSC lines were differentiated into ROs (Table S6). Samples were harvested at D140, D180 and/or D240. Overall, 3-6 replicates from 1-3 batches were collected. Batch quality was assessed by qPCR for photoreceptor-specific markers (Fig. S5). Only organoids exhibiting well-defined lamination and brush border morphology were selected for experiments. To generate iRPE cells, three control iPSC lines were used (Table S6). Samples were harvested at D98 (barely pigmented), D119 (pigmented but no stable TEER), and/or D140 (pigmented and stable TEER). Of note, differentiation days are calculated from the start of the differentiation, including the 3 passages that were performed. Overall, 5-6 replicates from 3 different batches were collected. Batch quality was assessed by TEER measurements and qPCR for RPE-specific markers (Fig. S6). Overall, all replicates and batches showed comparable differentiation rates in both ROs and iRPE. TEER measurements increased until ±D120, after which they reached a plateau until the day of collection.

### 4.2. TEER measurements

TEER measurements were taken using an EVOM3 Voltohmmeter (World Precision Instruments). Measurements were taken from at least 8 independent wells (24-well inserts) per iPSC-derived RPE line. Net TEER measurements were calculated by subtracting the value of a blank (geltrex-coated 24-well insert without cells) from the experimental value multiplied by the surface area (0.3 cm²).

### 4.3. POS supplementation to iRPE

Mature iRPE (i.e., D135) were treated for 4 days with bovine POS (41) and collected on day 5. On day 1, iRPE were pre-treated for 1h with 30% fetal bovine serum (FBS, sigma, F7524) to stimulate the phagocytosis process, after which approximately 5 POS per RPE cell were added combined with 10% FBS (27). On day 5, iRPE were washed 3 times with PBS to remove all non-ingested POS, and samples were collected using TrypLE (Gibco, 12604013).

### 4.4. Post-mortem human donor samples

Collection and sharing of *post-mortem* eyes was approved by the Medical ethics committee of Ghent University (UGent) and Ghent University Hospital (UZ Gent) (ONZ-2025-0105). Collection of the samples was conducted in accordance with the guidelines of the Declaration of Helsinki. In short, *post-mortem* residual ocular material obtained after cornea prelevation was collected from 6 donors by the Biobank Antwerpen (Antwerp, Belgium; ID: BE 71030031000) and the tissue bank of the Ghent University hospital. Samples were dissected within 24 hours *post-mortem*. The eye globe was dissected by removing the lens, iris, and vitreous, whereafter the choroid/RPE and neural retina were isolated. A small puncture (5 mm) was made in the neural retina, to collect samples from the macular. The remaining tissue was kept to analyze the non-macular regions. Following extraction, samples were washed 2 times with 0,9% NaCl, snap frozen in liquid nitrogen and stored at –80 °C until use.

### 4.5. RNA isolation, cDNA synthesis and qPCR

To assess the differentiation quality of the ROs and iRPE, a qPCR for differentiation markers was performed at D140 or D98, respectively. Two ROs were pooled, washed with PBS, snap frozen in liquid nitrogen and stored at –80 °C until use. Afterwards, ROs were submerged in RA1 buffer provided by the Nucleospin RNA kit (Macherey-Nagel, MN740955) and lysed using glass beads and the Tissue-LyserIII (Qiagen) at the frequency of 30/sec for 1 min. RPE cells were detached from the insert using TrypLE, pelleted at 1500 RPM for 5 min, washed 2 times with PBS, snap frozen in liquid nitrogen and stored in –80 °C until use. To lyse the RPE cells, RA1 buffer was added to the iRPE cells, and the sample was vortexed for 30 seconds. Next, RNA was extracted using the NucleoSpin RNA kit, converted to cDNA using IScript (Biorad, 1708891) and the qPCR was set-up using the GoTaq qPCR master mix (Promega, A6002), all following manufacturer’s instructions. Samples were run on a QuantStudio 5 Real-Time PCR system. To calculate the relative expression to a reference gene (*GUSB*), the 2^−ΔΔCT^ method was used. A full list of all the target genes and corresponding primer sequences is listed in Supplementary Table 7.

### 4.6. Untargeted lipidomics

For lipidomics, all RO, iRPE and *post-mortem* samples were washed 2 times with 0,9% NaCl (Merck, 1.06404.1000), flash frozen in liquid nitrogen and stored at –80 °C until use. iRPE cells were detached from the inserts using TrypLE. Overall, per replicate the following amounts were used: 6 ROs were pooled based on our pilot study (Swinkels *et al.* book chapter Retinal Degeneration, under review), RPE cells cultured on a 6-well insert, and half of a *post-mortem* human neural retina or RPE sample. Next, lipidome analysis was performed by the Core Facility Metabolomics of the Amsterdam UMC, as described previously (42). Lipid abundance was normalized to total protein concentration. Of note, only relative comparisons of the distribution of lipid species between samples could be performed due to sample heterogeneity, absolute quantitative comparisons were uninformative.

### 4.7. Statistics

The Metaboanalyst 6.0 software was used for the clustering analysis and fold-change graphs, while the other plots and graphs were made using Prism 10. Statistical analysis was performed using Prism 10 (GraphPad, San Diego, CA, USA). Grubbs’ test was executed on every dataset to identify possible outliers, the Shapiro–Wilk test was used to assess normal distribution, and the *F*-test was performed to test equality of the variances. Statistical significance was set at *P* < 0.05, and data are presented as mean ± SD. The statistical test used is indicated in the legend when applicable.

## 7. Acknowledgments

We would like to thank the Collin-Garanto lab for the discussions and feedback on this project. We also want to thank Eric Wever and Jan van Klinken from Amsterdam UMC for their bioinformatics support, Manon Huizing and Sarah Glorieux from Antwerp Biobank (Antwerp University Hospital, Antwerp, Belgium) and Tissue bank (Ghent University Hospital, Ghent, Belgium), respectively, as well as Dr. Anja Bufe from Boehringer Ingelheim. This work was funded by Stichting Oogfonds and Landelijke Stichting voor Blinden en Slechtzienden (UZ2023-17 and UZ2024-12 to DS, MAAP and AG) together with Stichting Blindenhulp, Rotterdamse Blindenbelangen, Het Lot and Gelderlandse Blindenstichting. It was also supported by NWO-XS (OCENW.XS24.3.048 to DS) and a Catalyst Grant from United for Metabolic Diseases (Metakids, 2023-CG-011 to DS, FMV and AG). Support was also obtained from the Radboudumc Therapy accelerator for rare diseases, financed by the Sectorplan Medical Sciences. EMvO, WK and ADMH are supported by Human Measurement Models 2.0 (grant nr. 18958) with additional funding from Proefdiervrij; Association of Collaborating Health Foundations (SGF), NWO Domain AES and the Netherlands Organisation for Health Research and Development (ZonMw), as part of their joint strategic research program: Human Measurement Models. The collaboration project is co-funded by the PPP Allowance made available by Health∼Holland, Top Sector Life Sciences & Health, to the SGF to stimulate public-private partnerships granted to AG. RJWC and AG were supported by the Ministry of Education, Culture and Science of the Netherlands (grant: Gravitation 024.006.034; Lifelong VISION. Finally, the work was supported by the Research Foundation Flanders (FWO) (1SD8924N to M.B. and G0ACQ24N to F.C.). The funding organizations had no role in the design or conduct of this research, and they provided unrestricted grants.

## 8. Limitations of the study

Retinal organoids and RPE require long and labor-intensive differentiation protocols. In this manuscript, we focused on three time points using our standard protocols. Future studies could investigate whether extended differentiation periods, lipid supplementation in the culture medium, or additional functional stimuli (e.g., light stimulation for retinal organoids or prolonged POS treatment) may alter the lipidome and make it more similar to that of *post-mortem* aged tissues.

Another important aspect is that our iPSC-derived models more closely resemble a newborn retina. However, we did not have access to embryonic tissue, which limited the comparisons that could be performed. At the same time, this limitation may provide insight into which lipids are required to induce aging in our *in vitro* models.

Finally, we performed untargeted lipidomics to obtain an overall overview of lipid classes across the different samples. To gain deeper insight into specific lipid classes or lipid species and to enable quantitative comparisons, more targeted approaches should be applied to accurately measure lipid content.

## 9. Resource availability

All lipidomics data generated in this study are available at the LIPID MAPS database. We are currently awaiting the DOI.

## 10. Author contributions

Conceptualization: DS and AG; Methodology: DS, FMV, AG; Scientific discussions: DS, EMVO, MB, ADMH, WK, FB, EDB, AB, SA, RWJC, FC, MAAPW, FMV, AG; Investigation: DS, EMVO, MB, ADMH, WK, FB; Funding acquisition: DS, FMV, MAAPW, AG,; Writing original draft: DS, AG; Review and editing: DS, EMVO, MB, ADMH, WK, FB, EDB, AB, SA, RWJC, FC, MAAPW, FMV, AG.

## 12. Declaration of generative AI and AI-assisted technologies

For some parts of the text, generative AI was used to reformulate sentences.

## References

1. Lewandowski D, Sander CL, Tworak A, Gao F, Xu Q, Skowronska-Krawczyk D. Dynamic lipid turnover in photoreceptors and retinal pigment epithelium throughout life. Prog Retin Eye Res. 2022;89:101037.

2. Swinkels D, Baes M. The essential role of docosahexaenoic acid and its derivatives for retinal integrity. Pharmacol Ther. 2023;247:108440.

3. Volland S, Esteve-Rudd J, Hoo J, Yee C, Williams DS. A comparison of some organizational characteristics of the mouse central retina and the human macula. PLoS One. 2015;10(4):e0125631.

4. Li B, Zhang T, Liu W, Wang Y, Xu R, Zeng S, et al. Metabolic Features of Mouse and Human Retinas: Rods versus Cones, Macula versus Periphery, Retina versus RPE. iScience. 2020;23(11):101672.

5. Gensheimer T, Veerman D, van Oosten EM, Segerink L, Garanto A, van der Meer AD. Retina-on-chip: engineering functional in vitro models of the human retina using organ-on-chip technology. Lab Chip. 2025;25(5):996–1014.

6. Rowe RG, Daley GQ. Induced pluripotent stem cells in disease modelling and drug discovery. Nat Rev Genet. 2019;20(7):377–88.

7. Zhang X, Wang W, Jin ZB. Retinal organoids as models for development and diseases. Cell Regen. 2021;10(1):33.

8. Afanasyeva TAV, Corral-Serrano JC, Garanto A, Roepman R, Cheetham ME, Collin RWJ. A look into retinal organoids: methods, analytical techniques, and applications. Cell Mol Life Sci. 2021;78(19-20):6505–32.

9. Kurzawa-Akanbi M, Tzoumas N, Corral-Serrano JC, Guarascio R, Steel DH, Cheetham ME, et al. Pluripotent stem cell-derived models of retinal disease: Elucidating pathogenesis, evaluating novel treatments, and estimating toxicity. Prog Retin Eye Res. 2024;100:101248.

10. Vasku G, Peltier C, He Z, Thuret G, Gain P, Gabrielle PH, et al. Comprehensive mass spectrometry lipidomics of human biofluids and ocular tissues. J Lipid Res. 2023;64(3):100343.

11. Anderson DMG, Messinger JD, Patterson NH, Rivera ES, Kotnala A, Spraggins JM, et al. Lipid Landscape of the Human Retina and Supporting Tissues Revealed by High-Resolution Imaging Mass Spectrometry. J Am Soc Mass Spectrom. 2020;31(12):2426–36.

12. Kotnala A, Anderson DMG, Messinger JD, Curcio CA, Schey KL. Untargeted Lipidomic Profiling of Aged Human Retina With and Without Age-Related Macular Degeneration (AMD). Adv Exp Med Biol. 2023;1415:37–42.

13. Bretillon L, Thuret G, Gregoire S, Acar N, Joffre C, Bron AM, et al. Lipid and fatty acid profile of the retina, retinal pigment epithelium/choroid, and the lacrimal gland, and associations with adipose tissue fatty acids in human subjects. Exp Eye Res. 2008;87(6):521–8.

14. Busik JV. Lipid metabolism dysregulation in diabetic retinopathy. J Lipid Res. 2021;62:100017.

15. Erion DM, Park HJ, Lee HY. The role of lipids in the pathogenesis and treatment of type 2 diabetes and associated co-morbidities. BMB Rep. 2016;49(3):139–48.

16. Anderson DMG, Ablonczy Z, Koutalos Y, Hanneken AM, Spraggins JM, Calcutt MW, et al. Bis(monoacylglycero)phosphate lipids in the retinal pigment epithelium implicate lysosomal/endosomal dysfunction in a model of Stargardt disease and human retinas. Sci Rep. 2017;7(1):17352.

17. Lakkaraju A, Umapathy A, Tan LX, Daniele L, Philp NJ, Boesze-Battaglia K, et al. The cell biology of the retinal pigment epithelium. Prog Retin Eye Res. 2020:100846.

18. Segawa K, Nagata S. An Apoptotic ‘Eat Me’ Signal: Phosphatidylserine Exposure. Trends Cell Biol. 2015;25(11):639–50.

19. Martin RE, Elliott MH, Brush RS, Anderson RE. Detailed characterization of the lipid composition of detergent-resistant membranes from photoreceptor rod outer segment membranes. Invest Ophthalmol Vis Sci. 2005;46(4):1147–54.

20. Agbaga MP, Merriman DK, Brush RS, Lydic TA, Conley SM, Naash MI, et al. Differential composition of DHA and very-long-chain PUFAs in rod and cone photoreceptors. J Lipid Res. 2018;59(9):1586–96.

21. Karjosukarso DW, Bukkems F, Duijkers L, Tomkiewicz TZ, Kiefmann J, Sarlea A, et al. Preclinical assessment of splicing modulation therapy for ABCA4 variant c.768G>T in Stargardt disease. Commun Med (Lond). 2025;5(1):25.

22. van Oosten EM, Hoogendoorn ADM, Kieboom W, Berendsen SL, Haerkens J, van der Maden MME, et al. Long-term daily light exposure boosts photoreceptor maturation in retinal organoids. bioRxiv. 2025.

23. Bose D, Ortolan D, Farnoodian M, Sharma R, Bharti K. Considerations for Developing an Autologous Induced Pluripotent Stem Cell (iPSC)-Derived Retinal Pigment Epithelium (RPE) Replacement Therapy. Cold Spring Harb Perspect Med. 2024;14(3).

24. Guymer R, Luthert P, Bird A. Changes in Bruch’s membrane and related structures with age. Prog Retin Eye Res. 1999;18(1):59–90.

25. Curcio CA, Presley JB, Malek G, Medeiros NE, Avery DV, Kruth HS. Esterified and unesterified cholesterol in drusen and basal deposits of eyes with age-related maculopathy. Exp Eye Res. 2005;81(6):731–41.

26. Neale BM, Fagerness J, Reynolds R, Sobrin L, Parker M, Raychaudhuri S, et al. Genome-wide association study of advanced age-related macular degeneration identifies a role of the hepatic lipase gene (LIPC). Proc Natl Acad Sci U S A. 2010;107(16):7395–400.

27. Mazzoni F, Mao Y, Finnemann SC. Advanced Analysis of Photoreceptor Outer Segment Phagocytosis by RPE Cells in Culture. Methods Mol Biol. 2019;1834:95–108.

28. West EL, Majumder P, Naeem A, Fernando M, O’Hara-Wright M, Lanning E, et al. Antioxidant and lipid supplementation improve the development of photoreceptor outer segments in pluripotent stem cell-derived retinal organoids. Stem Cell Reports. 2022;17(4):775–88.

29. Vainionpaa K, Kalatanova A, Seemab U, Montaser AB, Leinonen H. Female sex is a risk factor for exacerbated lipid peroxidation and disease in murine retinitis pigmentosa. Redox Biol. 2026;90:103987.

30. Hishikawa D, Valentine WJ, Iizuka-Hishikawa Y, Shindou H, Shimizu T. Metabolism and functions of docosahexaenoic acid-containing membrane glycerophospholipids. FEBS Lett. 2017;591(18):2730–44.

31. Walther TC, Farese RV, Jr. Lipid droplets and cellular lipid metabolism. Annu Rev Biochem. 2012;81:687–714.

32. Field FJ, Albright E, Mathur SN. Regulation of cholesterol esterification by micellar cholesterol in CaCo-2 cells. J Lipid Res. 1987;28(9):1057–66.

33. Rothblat GH, Arbogast LY, Ray EK. Stimulation of esterified cholesterol accumulation in tissue culture cells exposed to high density lipoproteins enriched in free cholesterol. J Lipid Res. 1978;19(3):350–8.

34. Raftopulos NL, Washaya TC, Niederprum A, Egert A, Hakeem-Sanni MF, Varney B, et al. Prostate cancer cell proliferation is influenced by LDL-cholesterol availability and cholesteryl ester turnover. Cancer Metab. 2022;10(1):1.

35. Boyer NP, Tang PH, Higbee D, Ablonczy Z, Crouch RK, Koutalos Y. Lipofuscin and A2E accumulate with age in the retinal pigment epithelium of Nrl−/− mice. Photochem Photobiol. 2012;88(6):1373–7.

36. Vives-Bauza C, Anand M, Shiraz AK, Magrane J, Gao J, Vollmer-Snarr HR, et al. The age lipid A2E and mitochondrial dysfunction synergistically impair phagocytosis by retinal pigment epithelial cells. J Biol Chem. 2008;283(36):24770–80.

37. Koolen L, Gagliardi G, Ten Brink SCA, de Breuk A, Heesterbeek TJ, Hoyng CB, et al. Generation and characterization of human induced pluripotent stem cells (iPSCs) from three individuals without age-related macular degeneration. Stem Cell Res. 2022;60:102670.

38. Karjosukarso DW, Bukkems F, Duijkers L, Leijsten N, Collin RWJ. Generation of three isogenic control lines from patient-derived iPSCs carrying bi-allelic ABCA4 variants underlying Stargardt disease. Stem Cell Res. 2023;71:103169.

39. Fligor CM, Huang KC, Lavekar SS, VanderWall KB, Meyer JS. Differentiation of retinal organoids from human pluripotent stem cells. Methods Cell Biol. 2020;159:279–302.

40. Regent F, Morizur L, Lesueur L, Habeler W, Plancheron A, Ben M’Barek K, et al. Automation of human pluripotent stem cell differentiation toward retinal pigment epithelial cells for large-scale productions. Sci Rep. 2019;9(1):10646.

41. ten Brink SCA, Koolen L, Klaver CCW, Bakker RA, den Hollander AI, Almedawar S. Non-canonical roles of CFH in retinal pigment epithelial cells revealed by dysfunctional rare CFH variants. Stem Cell Reports. 2025;20(1):102385.

42. Vaz FM, McDermott JH, Alders M, Wortmann SB, Kolker S, Pras-Raves ML, et al. Mutations in PCYT2 disrupt etherlipid biosynthesis and cause a complex hereditary spastic paraplegia. Brain. 2019;142(11):3382–97.

