## Supplementary figures and tables for "The Lipidome of iPSC-Derived Retinal Organoids and RPE Partially Resembles that of the Human Retina"

**Supplementary** **Tables**

*Supplementary table 1. Overview of the post-mortem human donor samples included in the lipidomics study.*

| Donor | Age (years) | Sex | Cause of death | Time to preservation (hours) |
| --- | --- | --- | --- | --- |
| 1 | 80 | Female | End stage COPD | 10 |
| 2 | 62 | Male | Cholangiocarcinoma | 12 |
| 3 | 80 | Male | Cancer | 22 |
| 4 | 68 | Male | Cardiopulmonary arrest | 20 |
| 5 | 58 | Female | Sarcoma | 22 |

*COPD-chronic obstructive pulmonary disease.*

*Supplementary Table 2. List of lipid species changed more than (-)5 fold in iRPE compared to pmRPE samples.*

*Supplementary Table 3. List of the 50 most abundant lipid species in the individual pmRPE samples and the three different iRPE lines.*

Green indicates lipid species that are in common between the different pmRPE samples. Red indicates those that are not in common.

*Supplementary Table 4. List of lipid species changed more than (-)5 fold in POS treated iRPE versus untreated iRPE or pmRPE samples.*

*Supplementary Table 5. List of lipid species changed more than (-)5 fold in post-mortem samples and/or ROs.*

*Supplementary table 6. Overview of the iPSC-derived RPE and RO samples included in the lipidomics study.*

| Published name | Label | Reprogramming method | Sex | Cell type | Harvest day | Batches | Replicates |
| --- | --- | --- | --- | --- | --- | --- | --- |
| HPSI0114i-kolf_3 | Male 1 | Sendai | Male | iRPE | D98  D119  D140 | 3  3  3 | 5  5  5 |
|  |  |  |  | RO | D140  D180  D240 | 3  3  3 | 5  5  5 |
| SCTCi048-A-1 | Male 2 | Episomal | Male | iRPE | D140 | 3 | 6 |
|  |  |  |  | RO | D240 | 2 | 5 |
| SCTCi010-A | Female 1 | Episomal | Female | iRPE | D98  D119  D140 | 3  3  3 | 6  6  6 |
|  |  |  |  | RO | D140  D180  D240 | 2  2  2 | 5  5  5 |
| RMCGENi005-A-1 | Female 2 | Lentivirus | Female | RO | D240 | 1 | 3 |

D-differentiation day; iRPE-iPSC-derived retinal pigment epithelium; RO-retinal organoid

*Supplementary table 7. Primer sequence list.*

| Primer name |  | Sequence (5’-3’) |
| --- | --- | --- |
| *CRX F* | RO marker | CCCCAGTGTGGATCTGATG |
| *CRX R* | RO marker | CAAACAGTGCCTCCAGCTC |
| *GUSB F* | Reference gene | AGAGTGGTGCTGAGGATTGG |
| *GUSB R* | Reference gene | CCCTCATGCTCTAGCGTGTC |
| *MERTK F* | RPE marker | TTGCAGCATTCAGGTCAAGGAAGC |
| *MERTK R* | RPE marker | GGCTTGCAGCTGCTTGATTTGGTA |
| *OCT4 F* | Pluripotency marker | GTTCTTCATTCACTAAGGAAGG |
| *OCT4 R* | Pluripotency marker | CAAGAGCATCATTGAACTTCAC |
| *OPSW F* | RO marker | TTCTTCTCCAAGAGTGCTTGC |
| *OPSW R* | RO marker | CCTTCCCACACACCATCTTC |
| *OPLW F* | RO marker | TCTGCTACCTCCAAGTGTGG |
| *OPLW R* | RO marker | CTTCCTTCTCTGCCTTCTGG |
| *RCVRN F* | RO marker | ACACCAAGTTCTCGGAGGAG |
| *RCVRN R* | RO marker | ACTTGGCGTAGATGCTCTGG |
| *RHO F* | RO marker | TCATCATGGTCATCGCTTTC |
| *RHO R* | RO marker | CATGAAGATGGGACCGAAGT |
| *RPE65 F* | RPE marker | TTACTACGCTTGCACAGAGACC |
| *RPE65 R* | RPE marker | GCCCCATTGACAGAGACATAG |
| *VMD F* | RPE marker | TCAGTGTGGACACCTGTATGC |
| *VMD R* | RPE marker | AAGCTGTACACCGCCACAG |

**Supplementary figures**

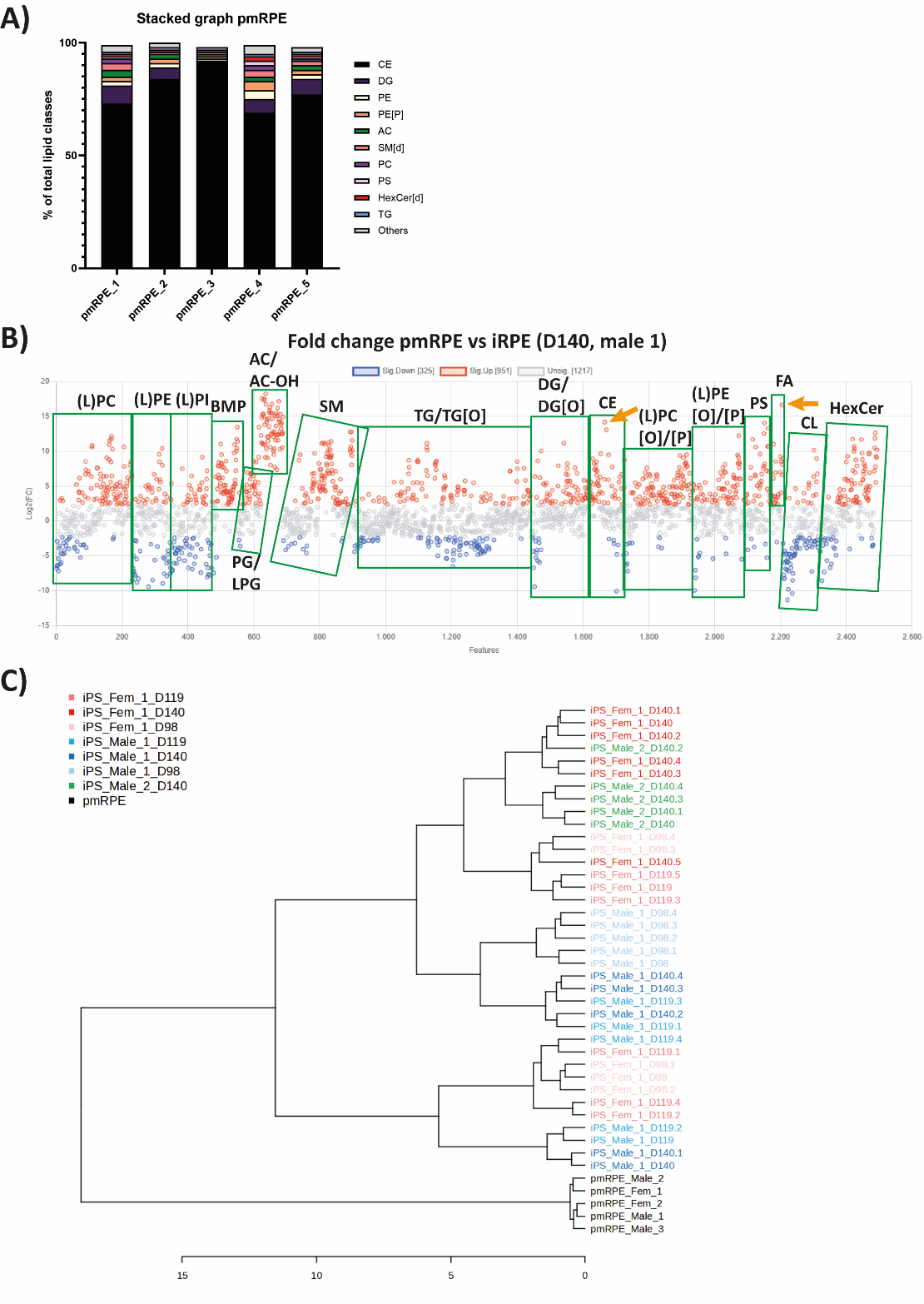

***Supplementary Figure 1. Differential lipid composition in iRPE compared to pmRPE.*** *a) The relative contribution of different lipid classes to the total lipidome in pmRPE samples. b) Graph representing the fold-change of lipid species in pmRPE versus iRPE (D140, male 1). Treshhold was set at FC of (-)5. Orange arrow indicates lipids containing DHA (C22:6). c) Hierarchical clustering dendogram of pmRPE and different time-points and control lines of iRPE using the Pearson Ward test. (L)PC-(lyso-)phospatidylcholine; (L)PC[O]-(lyso)alkyllysophosphatidylcholine; (L)PC[P]-(lyso)alkenyllysophosphatidylcholine; (L)PE-(lyso)phosphatidylethanolamine; (L)PE[O]-(lyso)alkyllysophosphatidylethanolamine; (L)PE[P]-(lyso)alkyllysophosphatidylethanolamine; (L)PG-(lyso)phosphatidylglycerol; (L)PI-(lyso)phosphatidylinositol; AC-acylcarnitines; AC-OH-hydroxy acylcarnitine; BMP-bis(monoacylglycerol)phosphate; CE-cholesteryl esters; Cer[d]-ceramide; CL-cardiolipin; D-differentiation day; DG[O]-alkylacylglycerol; DG-diacylglycerol; FA-fatty acid; HexCer[d]-hexosylceramides; PE[P]-alkenyl phosphatidylethanolamine; PS-phosphatidylserine; SM[d]-sphingomyelin; TG[O]-alkyldiacylglycerol; TG-triglycerides.*

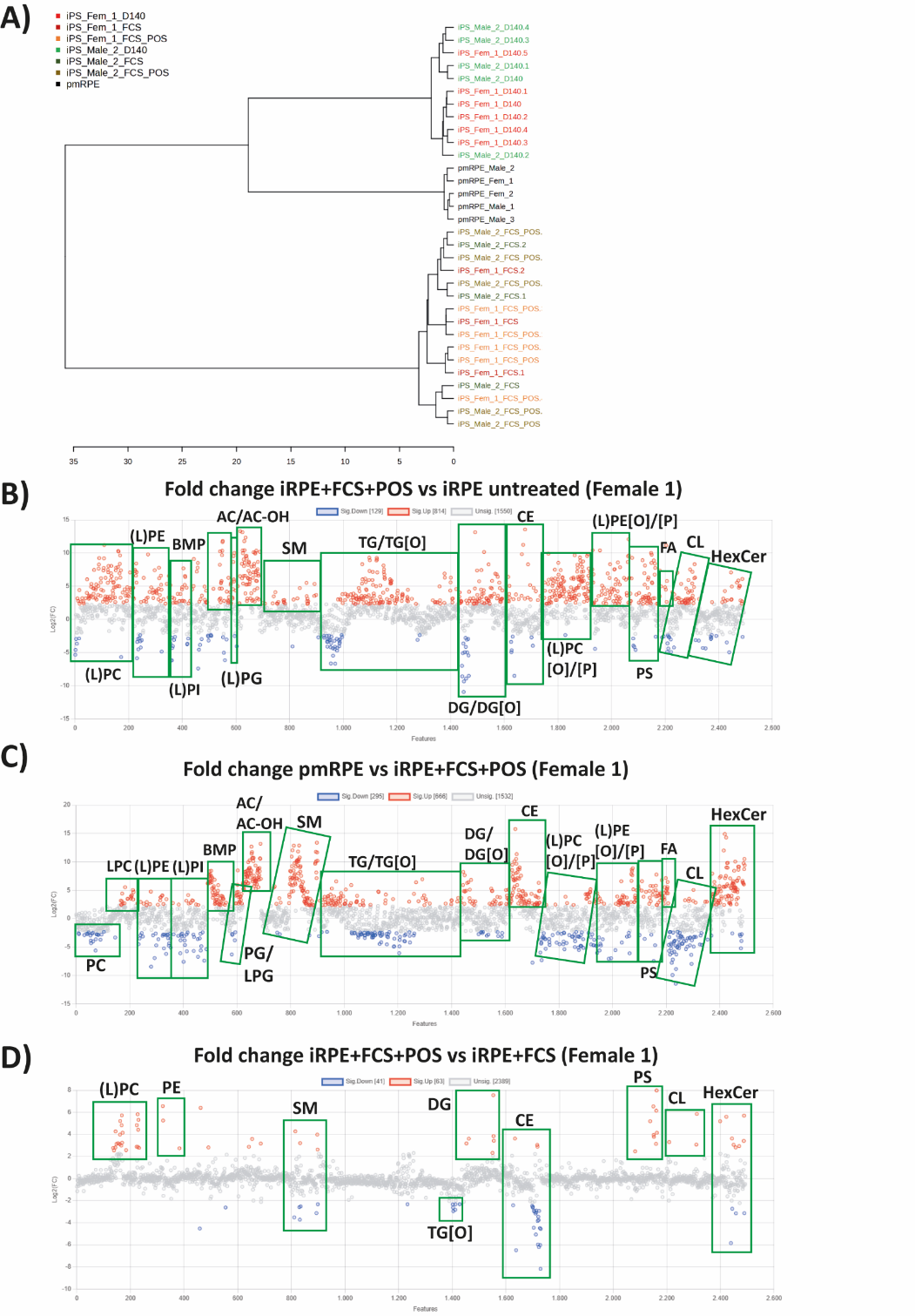

***Supplementary Figure 2. Differential lipid composition in pmRPE compared to POS-fed iRPE.*** *a) Hierarchical clustering dendogram on pmRPE and different control lines of iRPE not treated, FCS-treated or FCS+POS using the Pearson Ward test. b) Graph representing the fold-change of lipid species in untreated iRPE (D140, female 1) versus FCS+POS treated iRPE (D140, female 1). c) Graph representing the fold-change of lipid species in pmRPE (average) versus FCS+POS treated iRPE (D140, female 1). d) Graph representing the fold-change of lipid species in FCS+POS treated iRPE (D140, female 1) versus FCS treated iRPE (D140, female 1). Treshhold was set at FC of (-)5. (L)PC-(lyso-)phospatidylcholine; (L)PC[O]-(lyso)alkyllysophosphatidylcholine; (L)PC[P]-(lyso)alkenyllysophosphatidylcholine; (L)PE-(lyso)phosphatidylethanolamine; (L)PE[O]-(lyso)alkyllysophosphatidylethanolamine; (L)PE[P]-(lyso)alkyllysophosphatidylethanolamine; (L)PG-(lyso)phosphatidylglycerol; (L)PI-(lyso)phosphatidylinositol; AC-acylcarnitines; AC-OH-hydroxy acylcarnitine; BMP-bis(monoacylglycerol)phosphate; CE-cholesteryl esters; Cer[d]-ceramide; CL-cardiolipin; D-differentiation day; DG[O]-alkylacylglycerol; DG-diacylglycerol; FA-fatty acid; HexCer[d]-hexosylceramides; PE[P]-alkenyl phosphatidylethanolamine; PS-phosphatidylserine; SM[d]-sphingomyelin; TG[O]-alkyldiacylglycerol; TG-triglycerides.*

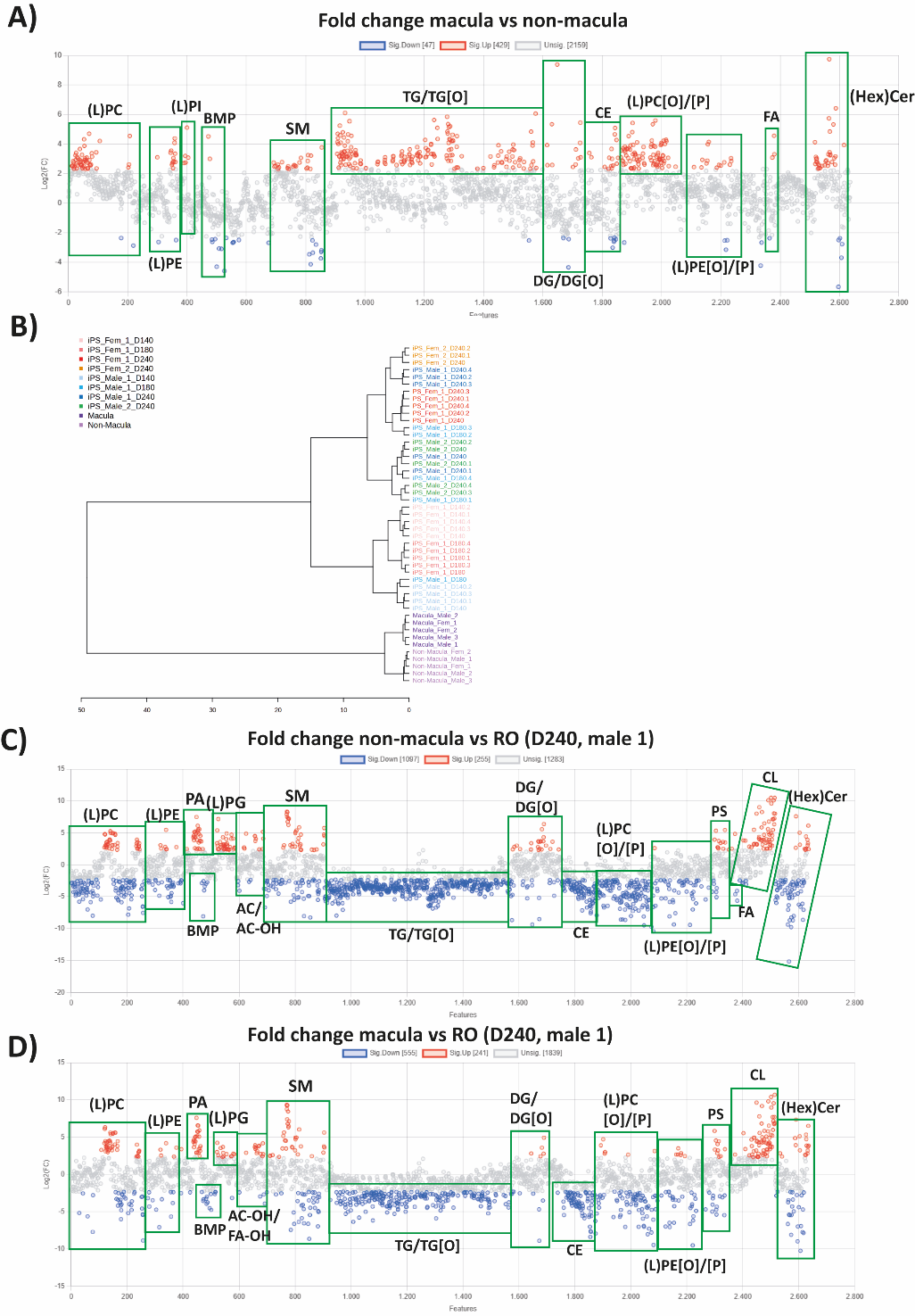

***Supplementary Figure 3. Differential lipid composition post-mortem macula and non-macula and ROs.*** *a) Graph representing the fold-change of lipid species in post-mortem macula versus non-macula samples. b) Hierarchical clustering dendogram on post-mortem macula, non-macula and different control lines and time-points of ROs using the Pearson Ward test. c) Ratio of Ꞷ3/Ꞷ6 lipid species in post-mortem and RO samples. Ꞷ3-species included LPC, LPE, LPI, LPA, LPG, LPS and FA-containing C22:6. Ꞷ6-species included LPC, LPE, LPI, LPA, LPG, LPS and FA-containing C20:4.* *d-e) Graph representing the fold-change of lipid species in post-mortem non-macula (c) or macula (d) versus RO (D240, male 1). Treshhold was set at FC of (-)5. (L)PC-(lyso-)phospatidylcholine; (L)PC[O]-(lyso)alkyllysophosphatidylcholine; (L)PC[P]-(lyso)alkenyllysophosphatidylcholine; (L)PE-(lyso)phosphatidylethanolamine; (L)PE[O]-(lyso)alkyllysophosphatidylethanolamine; (L)PE[P]-(lyso)alkyllysophosphatidylethanolamine; (L)PG-(lyso)phosphatidylglycerol; (L)PI-(lyso)phosphatidylinositol; AC-acylcarnitines; AC-OH-hydroxy acylcarnitine; BMP-bis(monoacylglycerol)phosphate; CE-cholesteryl esters; Cer[d]-ceramide; CL-cardiolipin; D-differentiation day; DG[O]-alkylacylglycerol; DG-diacylglycerol; FA-fatty acid; HexCer[d]-hexosylceramides; PE[P]-alkenyl phosphatidylethanolamine; PS-phosphatidylserine; SM[d]-sphingomyelin; TG[O]-alkyldiacylglycerol; TG-triglycerides.*

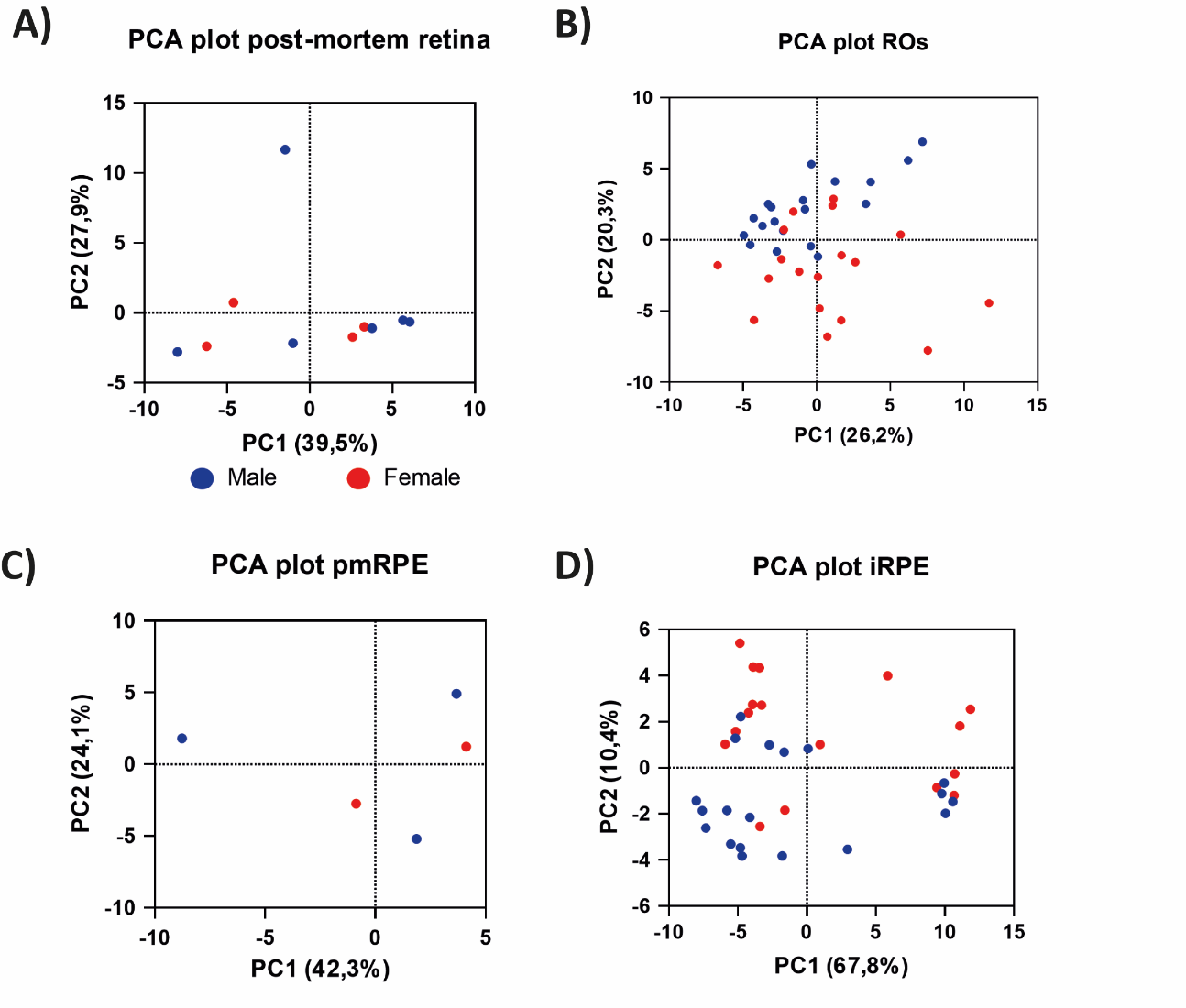
***Supplementary Figure 4. No substantial differences in lipid profile between male and female retina.*** *a-d) PCA plots showing replicates from post-mortem retina (a), ROs (b), pmRPE (c) and iRPE (d) clustered according to their variance in two dimensions.*

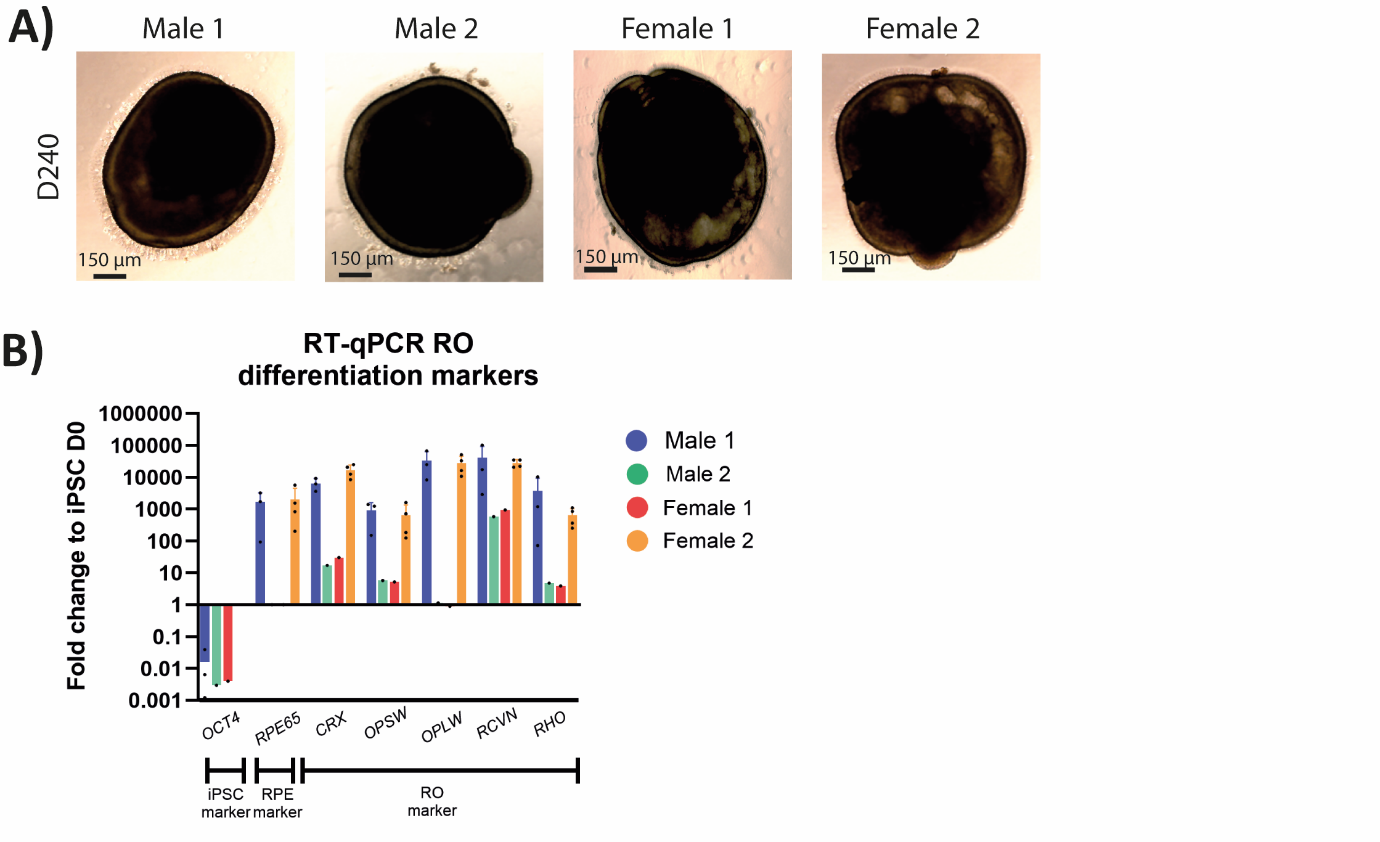

***Supplementary figure 5. Differentiation quality assessment ROs.*** *a) Representative bright field images of the four different control iPSC-derived ROs at D240 used in this study. ROs with clear lamination and brush border morphology were selected for the study. b) RT-qPCR for iPSC (OCT4, negative control), RPE (RPE65, negative control) and photoreceptor markers (CRX, OPSW, OPLW, RCVRN, RHO). differentiation markers. Data are expressed as fold-change to Day 0 of the differentiation.*

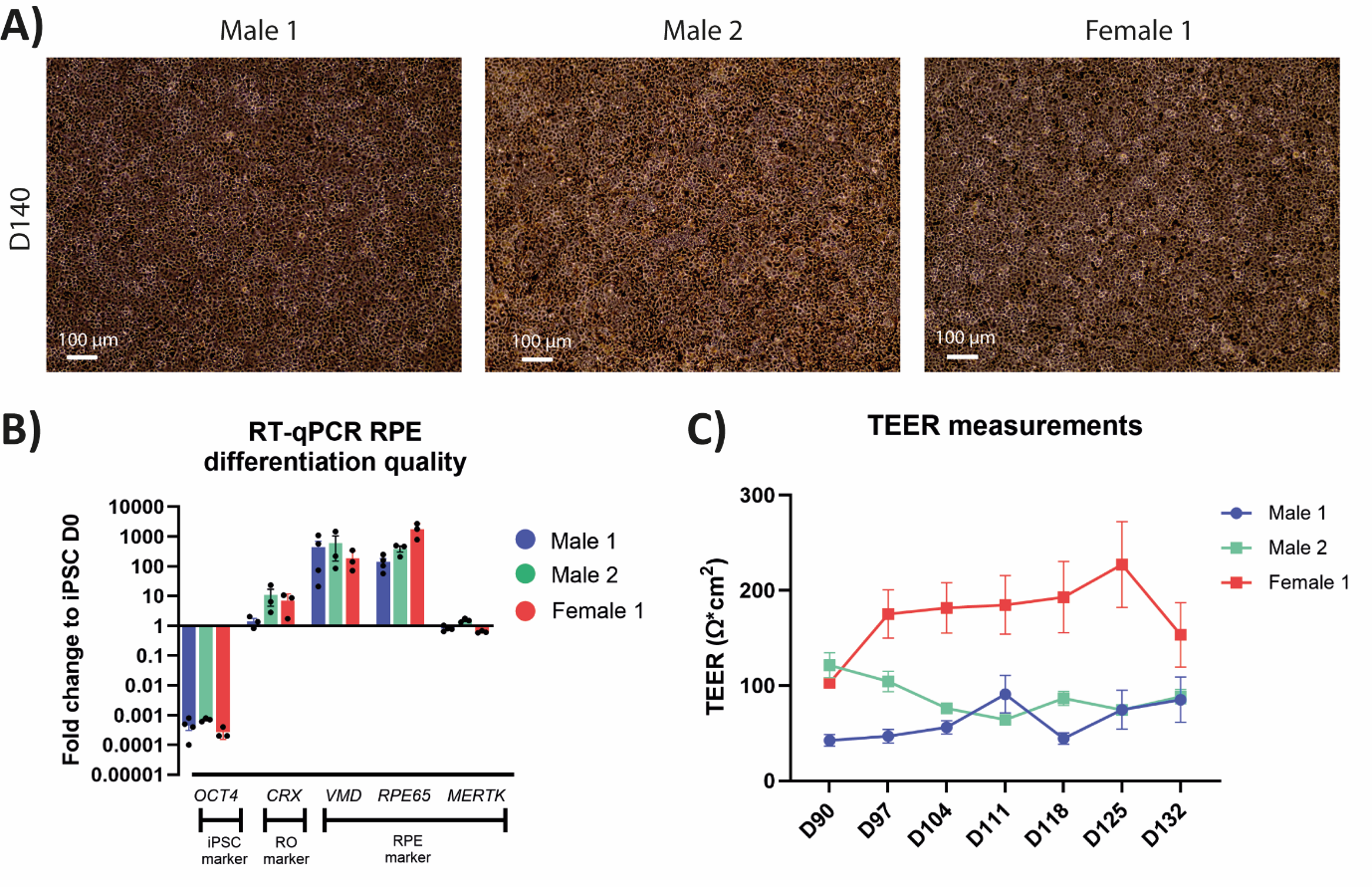
***Supplementary figure 6. Differentiation quality assessment iRPE.*** *a) Representative bright field images of the three different control iPSC-derived RPE at D140 used in this study. iRPE with clear honeycomb organization and pigmentation were selected for the study. b) RT-qPCR for iPSC (OCT4, negative control), RO (CRX, negative control) and RPE (BEST1, RPE65, MERKT) markers. Data are expressed as fold-change to Day 0 of the differentiation. c) TEER (transepithelial electrical resistance) measurements expressed in Ω, showing a gradual increase in resistance during the differentiation process.*
